# BK polyomavirus (BKPyV) is a risk factor for bladder cancer through induction of APOBEC3-mediated genomic damage

**DOI:** 10.1101/2021.05.13.443803

**Authors:** Simon C. Baker, Andrew S. Mason, Raphael G. Slip, Katie T. Skinner, Andrew Macdonald, Omar Masood, Reuben S. Harris, Tim R. Fenton, Manikandan Periyasamy, Simak Ali, Jennifer Southgate

## Abstract

Limited understanding of bladder cancer aetiopathology hampers progress in reducing incidence. BK polyomavirus (BKPyV) is a common childhood infection that can be reactivated in the adult kidney leading to viruria. Here we used a mitotically-quiescent, differentiated, normal human urothelial *in vitro* model to study BKPyV infection. BKPyV infection led to significantly elevated APOBEC3A and APOBEC3B protein, increased deaminase activity and greater numbers of apurinic/apyrimidinic sites in the host urothelial genome. BKPyV Large T antigen (LT-Ag) stimulated re-entry into the cell cycle via inhibition of Retinoblastoma protein and activation of EZH2, E2F1 and FOXM1, which combined to push urothelial cells from G0 into an arrested G2 cell cycle state. The single-stranded DNA displacement loops formed during BKPyV-infection, provide a substrate for APOBEC3 enzymes where they interacted with LT-Ag. These results support reactivated BKPyV infections in adults as a risk factor for bladder cancer in immune-insufficient populations, including transplant patients and the elderly.

## Introduction

Urothelial (bladder) cancer has a complex natural history with an indolent, frequently recurrent and unpredictably progressive disease path, which converges with a more aggressive route taken by malignancies that can present at advanced muscle-invasive and even disseminated stages. Although smoking is a well-established risk factor for bladder cancer (BLCA), the mutational signatures of bladder tumours show only a minor proportion of the G>T transversions characteristic of DNA damage caused directly by smoke-derived carcinogens^1^. Polyomavirus (PyV) infection could provide an alternative route to initiation of particular relevance to immune-insufficient populations.

PyV are ubiquitous childhood infections, with 70-80% seroprevalence by adulthood^2^. They are largely asymptomatic in the immuno-competent, but may remain latent in the adult kidney^3^. BK polyomavirus (BKPyV) resides latent in differentiated renal proximal tubule cells and reactivation at times of immune-insufficiency leads to sloughing of actively infected “decoy” cells (into the urine), to limit kidney damage^4^. PyV DNA can be detected in the urine of immuno-competent individuals with a frequency that increases with age (to ≥30% in the over 50s for BKPyV)^5^. The only published study of BLCA risk (n=3,782), found a higher incidence (15.8%) in the 133 patients with previous urine cytology evidence of BKPyV infection (OR3.4, p < 0.001)^6^.

Studies of bladder tumour genomes have identified mutational signatures associated with the anti-viral apolipoprotein B mRNA editing enzyme catalytic polypeptide (APOBEC) family of cytosine deaminase enzymes in up to 93% of cases^7^. The large T antigen (LT-Ag) of BKPyV has been shown to induce APOBEC3B expression^8, 9^. However, studies of bladder tumours fail to identify viral DNA or RNA, with fewer than 4% positives reported in the largest study of 689 cases^10^. A successful PyV life-cycle requires the viral genome to remain episomal (*i.e.* non-integrated). The hypothesis of this study is that the episomal life-cycle of BKPyV in the urothelium is capable of initiating bladder tumours via APOBEC-mediated damage of the host genome. This is a so-called “hit-and-run” mode of carcinogenesis whereby the presence of virus causes the initiating inactivation mutation of tumour suppressor genes decades before a tumour forms. Subsequent immune clearance of BKPyV leads to its absence from later stages of tumour development (summarised Fig. 1).

**Fig. 1.**
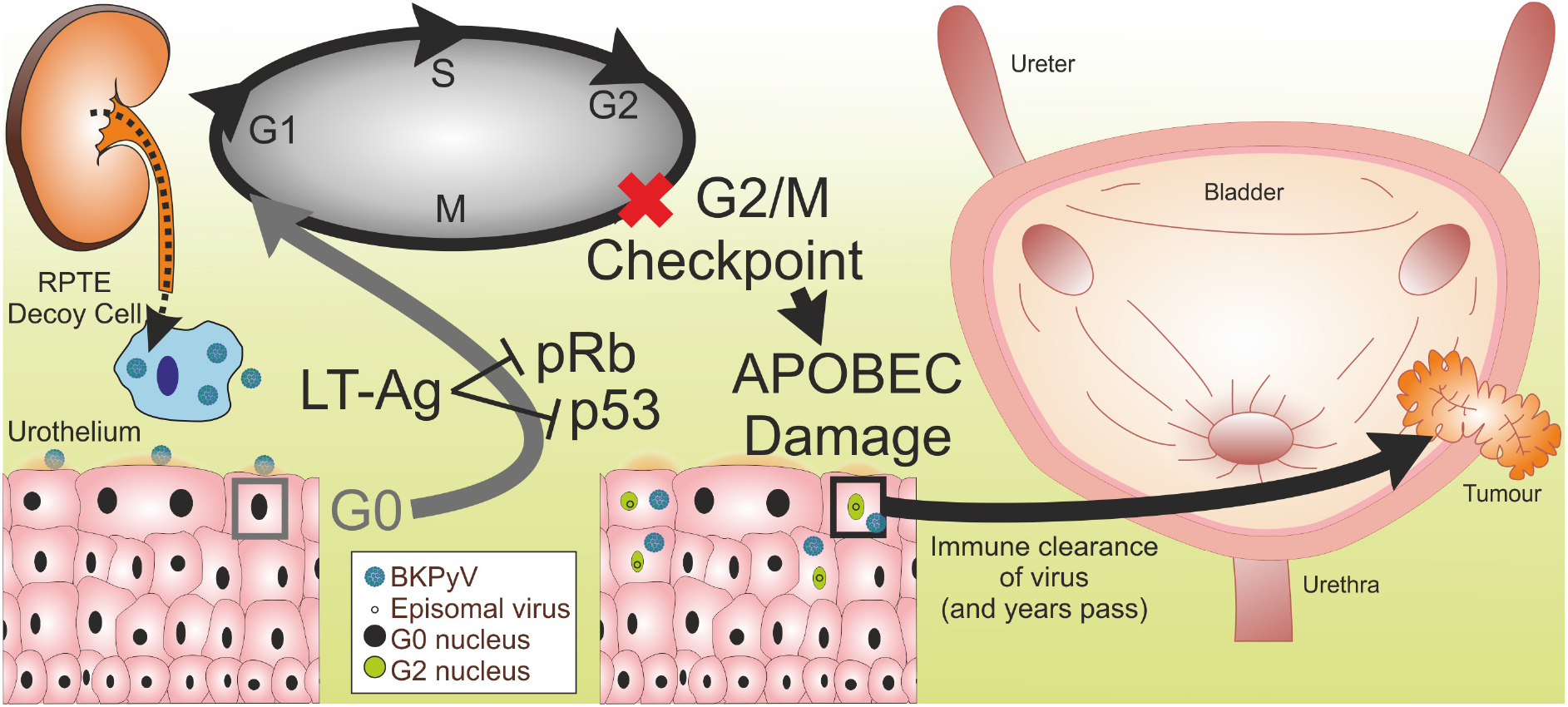
Schematic model of BKPyV hit-and-run carcinogenesis hypothesis. Immune-insufficiency leads to sloughing of actively infected renal “decoy” cells and BKPyV viruria. BKPyV infects the G0-arrested urothelium. BKPyV LT-Ag inhibits retinoblastoma (pRb) and p53 inhibition of the cell cycle. BKPyV cell cycle re-entry progresses to arrest at the G2/M checkpoint. BKPyV stimulates APOBEC3 enzyme activity and host genome damage that inactivates tumour suppressors. The immune system clears the virus but initiated cells persist and over a period of years expand to form a tumour.

Due to the asymptomatic nature of BKPyV infection, clinical sampling during reactivated adult infection is challenging and since BKPyV is human-specific, *in vivo* models are not applicable. This study was designed to evaluate hit-and-run carcinogenic mechanisms for BKPyV-infection of the normal human urothelium *in vitro*. Human urothelium is a low-turnover mitotically-quiescent epithelium where cells reside in G0^11^. To address limitations in previous urothelial studies which have employed actively-dividing, undifferentiated cell cultures^12–14^; this study applied a unique, quiescent (G0-arrested), stratified and barrier-forming *in vitro* model of normal human urothelium that uses biological replicates to reflect donor diversity^15^. Interferon-γ (IFNγ) has previously been clinically associated with BKPyV infection^16^ and suggested to reduce BKPyV-infection progression in renal cell cultures^17^; therefore we investigated its potential for regulating urothelial anti-viral self-defence mechanisms.

## Results

### BKPyV Infection of normal human urothelium

Initial studies found BKPyV infection of mitotically-quiescent urothelium to be more successful in differentiated than undifferentiated cultures (Extended Data Fig. 1). For this study, BKPyV infection was evaluated during the pre-lytic phase of infection in differentiated normal human urothelial (NHU) cell cultures which showed no morphological or transcriptomic signs of apoptosis during the 14 day infection period (Extended Data Fig. 2).

**Fig. 2.**
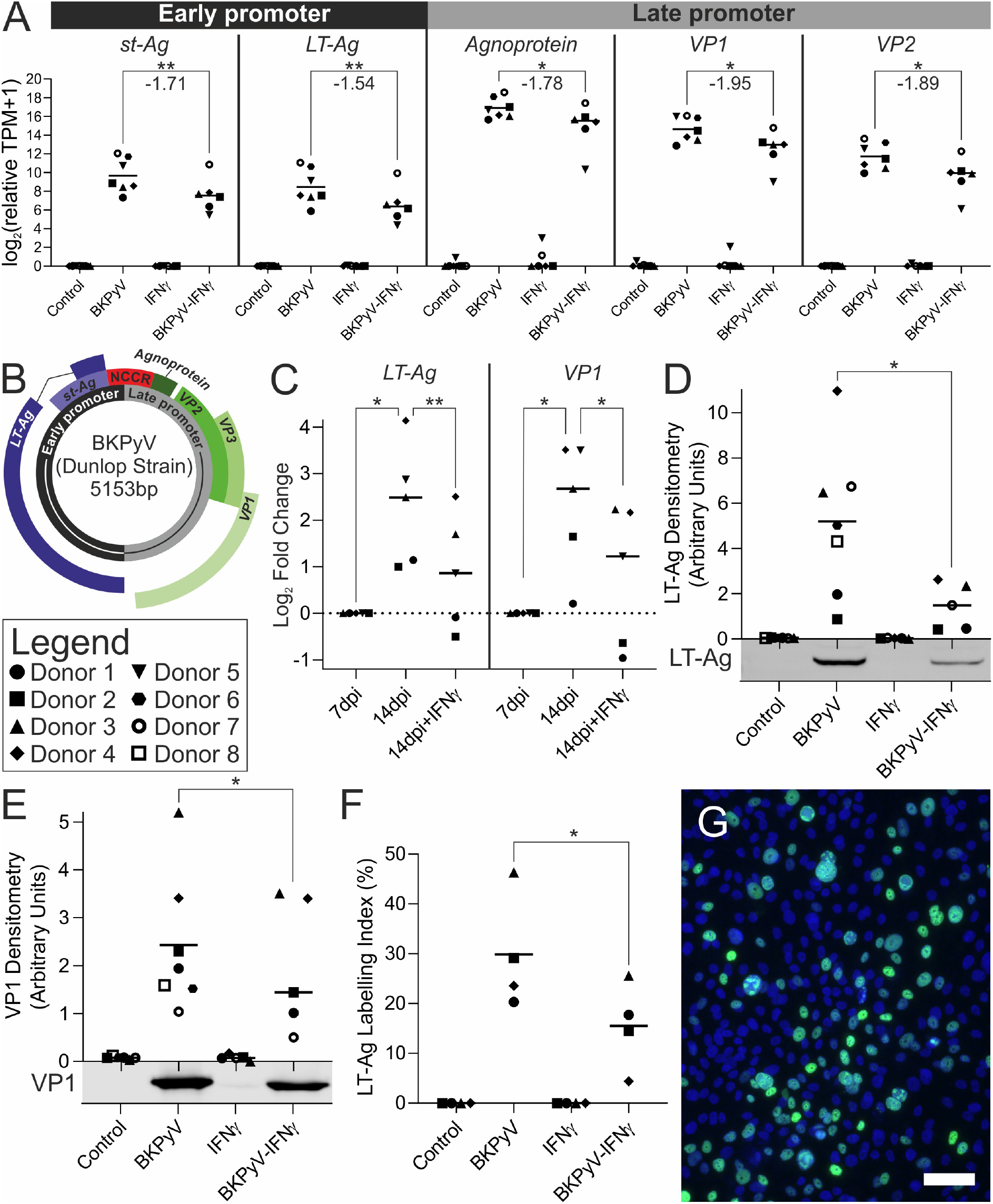
(A) mRNAseq analysis of BKPyV gene expression at 14dpi shows Agnoprotein was the most expressed viral gene and LT-Ag the least expressed. All viral gene expression was significantly suppressed by IFNγ; however, the efficacy was widely variable between donors. Statistically significant comparisons are indicated by stars with the mean log2 fold change in gene expression reported beneath (n=6/7 independent donors). (B) BKPyV genome map showing the non-coding control region (NCCR) which regulates both the early and late genes that are expressed in opposing orientations. (C) RT-qPCR analysis of BKPyV LT-Ag and VP1 transcript abundance in NHU cell cultures. Data is displayed as log2 fold-change normalised to abundance at 7 days post infection (dpi). VP1 abundance increased significantly between 7 and 14dpi. The addition of IFNγ significantly reduced VP1 abundance at 14dpi. (D) 14dpi Western blot densitometry for large T antigen (LT-Ag) with exemplar blot image below the X-axis. Truncated LT-Ag (truncT-Ag) was also expressed; densitometry analysis of truncT-Ag and whole blots for LT-Ag can be found in Extended Data Fig. 3 along with the β-actin loading controls for all blots in Extended Data Fig. 4. (E) 14dpi Western blot densitometry for viral capsid protein 1 (VP1) with exemplar blot image below x-axis. Full blots are shown as Extended Data Fig. 5 (n=5 independent donors). (F) LT-Ag indirect immunofluorescence labelling index. n>2900 cells per condition per donor. (G) Exemplar LT-Ag indirect immunofluorescence image from Donor 3 BKPyV-infected urothelial cells (Scale bar denotes 10 μm). Significance was assessed in panels A by LRT test and B-E by paired t-test.

This is the first transcriptomic study of BKPyV infection of normal human urothelial tissues, which reflect human diversity in utilising cultures from multiple different donors (n=7 for mRNAseq). All genes of the BKPyV genome were expressed at 14dpi, with *Agnoprotein* the most (mean relative TPM=157,998) and *LT-Ag* the least (mean relative TPM=699) expressed transcripts (Fig. 2A with genome map as Fig. 2B). Overall, the expression pattern is indicative of greater activity at the late promoter driving *Agnoprotein*, *VP1* and *VP2* gene expression (Fig. 2A). In some cultures, IFNγ was added for the 7-14dpi period and the cytokine significantly reduced expression of all viral genes to an average of 44% of that observed in the absence of IFNγ (Fig. 2A). However, the ability of human urothelium to frustrate BKPyV gene expression in the presence of IFNγ was highly variable between donors, with the IFNγ-induced reduction in expression ranging from 1.8% to 66.2% in donors 5 and 3 respectively (Fig. 2A).

RT-qPCR was employed to evaluate *LT-Ag* and *VP1* transcript expression during the development of infection from 7-14dpi (Fig. 2C). In all infected cultures, viral gene expression increased from 7dpi to 14dpi (Fig. 2C). In cultures from donors 1&2, the addition of IFNγ at 7dpi reduced the *LT-Ag* and *VP1* transcript burden by 14dpi, whereas in the remaining three donors tested, the increase in *LT-Ag* and *VP1* transcripts was merely attenuated (Fig. 2C).

Western blotting for viral proteins LT-Ag (including detection of Truncated LT-Ag “truncT-Ag”^18^; Extended Data Fig. 3) and VP1 found both were significantly (*p*<0.05) reduced by addition of IFNγ (Fig. 2D&E). Indirect immunofluorescence labelling of LT-Ag showed that the addition of IFNγ led to significantly fewer infected cells being detected in cultures at 14dpi (Fig. 2F&G).

**Fig. 3.**
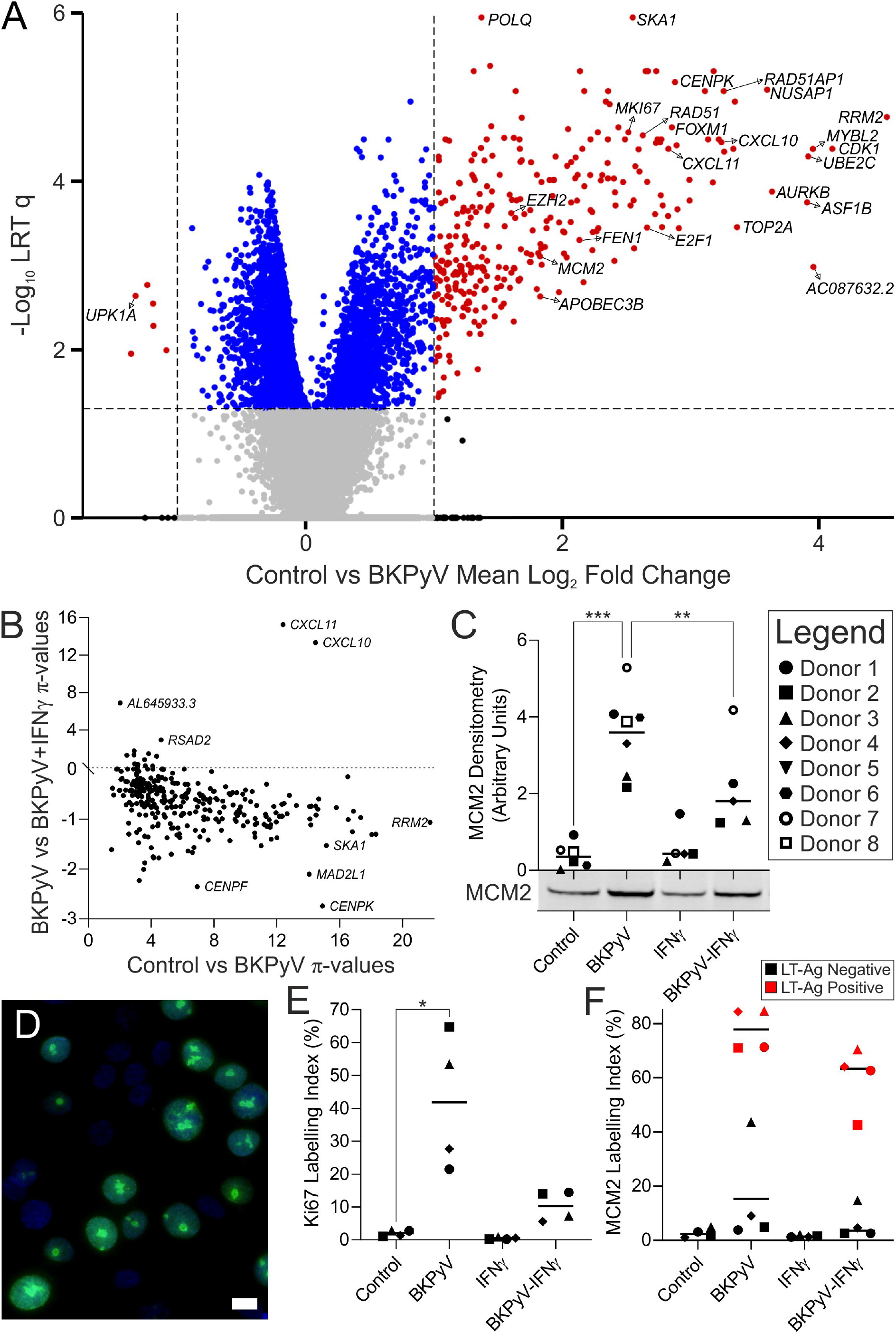
mRNAseq analysis of BKPyV infection of differentiated human urothelium. (A) Volcano plot highlighting the significant induction of cell cycle and DNA damage genes during BKPyV infection compared to controls (n=7 independent donors). (B) The 305 genes significantly induced by BKPyV-infection were plotted to show their induction by BKPyV in relation to the post-infection addition of IFNγ. Addition of IFNγ suppressed expression of the vast majority of BKPyV-induced genes (t=−17.57; p=1.97×10^−47^). The chemokines CXCL10 and CXCL11 were notable exceptions, where the addition of IFNγ dramatically increased expression. (C) 14dpi Western blot densitometry for DNA replication licensing factor MCM2 with exemplar blot image below x-axis. Whole blots for MCM2 can be found in Extended Data Fig. 8. (D) Indirect immunofluorescence labelling of Ki67 (green) in BKPyV infected NHU cells shows nearly all Ki67-positive cells have a few large nucleolar granules, characteristic of the G2 cell cycle stage. DNA was stained with Hoechst 33258 (blue). White scale bar denotes 10 μm. (E) Ki67 labelling indices for quiescent, G0-arrested control cultures were low (mean=1.99% ±0.94). BKPyV infection led to a significant (p=0.0319) increase in Ki67 labelling index (mean=41.8 ±20.62). (F) BKPyV infection also led to a significant increase in MCM2 labelling index (p<0.05; Extended Data Fig. 10). When cells from infected cultures were separated into LT-Ag positive and negative populations, there was a mean 5.6-fold (±4.0) increase in MCM2 labelling of LT-Ag negative cells from BKPyV-infected cultures from all donors when compared to controls. The red colouration for LT-Ag positive cells applies only to panel F.

Analysis of the human urothelial transcriptome at 14dpi with BKPyV by mRNAseq found 305 transcripts were *q*<0.05 significantly >2-fold induced (Fig. 3A). The major task of any PyV infection of the urinary tract is the initiation of proliferation, since the target epithelia (namely proximal tubule and urothelium) are not actively dividing tissues and both are predominantly (>99%) out-of-cycle, arrested in G0. Gene set enrichment analysis (GSEA; Extended Data Table 1) suggested transcriptional programmes related to cell cycle re-entry (specifically the G2/M checkpoint) and DNA damage repair were activated in the mitotically quiescent urothelial cells (Extended Data Fig. 6A&B). A small number of genes were significantly reduced; these included the uroplakin genes *UPK1A*, *UPK2* and *UPK3A*, suggesting superficial urothelial differentiation was affected by BKPyV infection (Fig. 3A). Analysis of the BKPyV-induced urothelial transcriptome showed IFNγ significantly suppressed expression of the 305 infection-induced genes (t=−17.57; *p*=1.97×10^−47^; Fig. 3B).

Re-entry into the cell cycle was supported by significant (*q*<0.001) *MCM2* and *MKI67* transcript induction (mean log2 fold change = 5.69 and 2.11, respectively; Extended Data Fig. 6C&D). IFNγ exposure reduced mean *MCM2* and *MKI67* transcript expression in BKPyV-infected cultures compared with infection alone but, due to the variance in the IFNγ-response, these changes were not statistically significant (Extended Data Fig. 6C&D). At the protein level MCM2 was significantly induced by BKPyV infection (*p*<0.001) and that increase was significantly suppressed by IFNγ (*p*<0.01; Fig. 3C & Extended Data Fig. 7). Cell cycle stage analysis of Ki67 immunolocalisation^19^ showed both a significant (*p*=0.03) increase in the number of positive cells within BKPyV infected cultures and that Ki67-positive cells were overwhelmingly in the G2 stage of the cell cycle, as identified by the few large nucleolar granules observed in each nucleus (Fig. 3D and E, respectively; Extended Data Fig. 8). G2 arrest in BKPyV infected cultures was supported transcriptomically by significant enrichment of gene sets associated with negative regulation of the G2 to M transition and experimentally-induced G2-arrest by 2-methoxyestradiol (Extended Data Fig. 6E&F). Furthermore, analysis of nuclei in cells from BKPyV-infected cultures indicated a significant increase in nuclear size for LT-Ag labelled cells, supporting cell cycle progression beyond S-phase (Extended Data Fig. 9). Indirect immunofluorescence co-labelling of MCM2 and the LT-Ag revealed that BKPyV triggered a significant increase in MCM2 positive cells and that significantly fewer cells became MCM2 positive in the presence of IFNγ (Extended Data Fig. 10). Using LT-Ag labelling to differentiate positive from negative cells in infected cultures indicated that MCM2 was also elevated 5.6-fold (±4.0) in LT-Ag negative cells within BKPyV-exposed cultures (Fig. 3F).

BKPyV regulates the cell cycle in part via LT-Ag interactions through its L×C×E domain with Retinoblastoma protein (pRb; Fig. 4A). GSEA supported the role of pRb disruption (Extended Data Fig. 12A-C) in driving proliferation, and phosphorylation of pRb was significantly (*p*<0.05) both increased by BKPyV and reduced by IFNγ exposure (Fig. 4B; Extended Data Fig. 13). Indirect immunofluorescence co-labelling of phosphorylated-pRb and the LT-Ag revealed BKPyV triggered an increase in phosphorylated-pRb labelled cells and that significantly fewer cells became positive in the presence of IFNγ (both *p*<0.01; Extended Data Fig. 11). Using LT-Ag labelling (to distinguish infected from non-infected cells in infected cultures), 63.1-fold (±57.1) more phosphorylated-pRb positive cells were present in the LT-Ag negative fraction of BKPyV-exposed cultures compared to control cultures (Fig. 4C). pRb phosphorylation and cell cycle re-entry by infected and adjacent cells could contribute to tumour promotion in initiated cells.

**Fig. 4.**
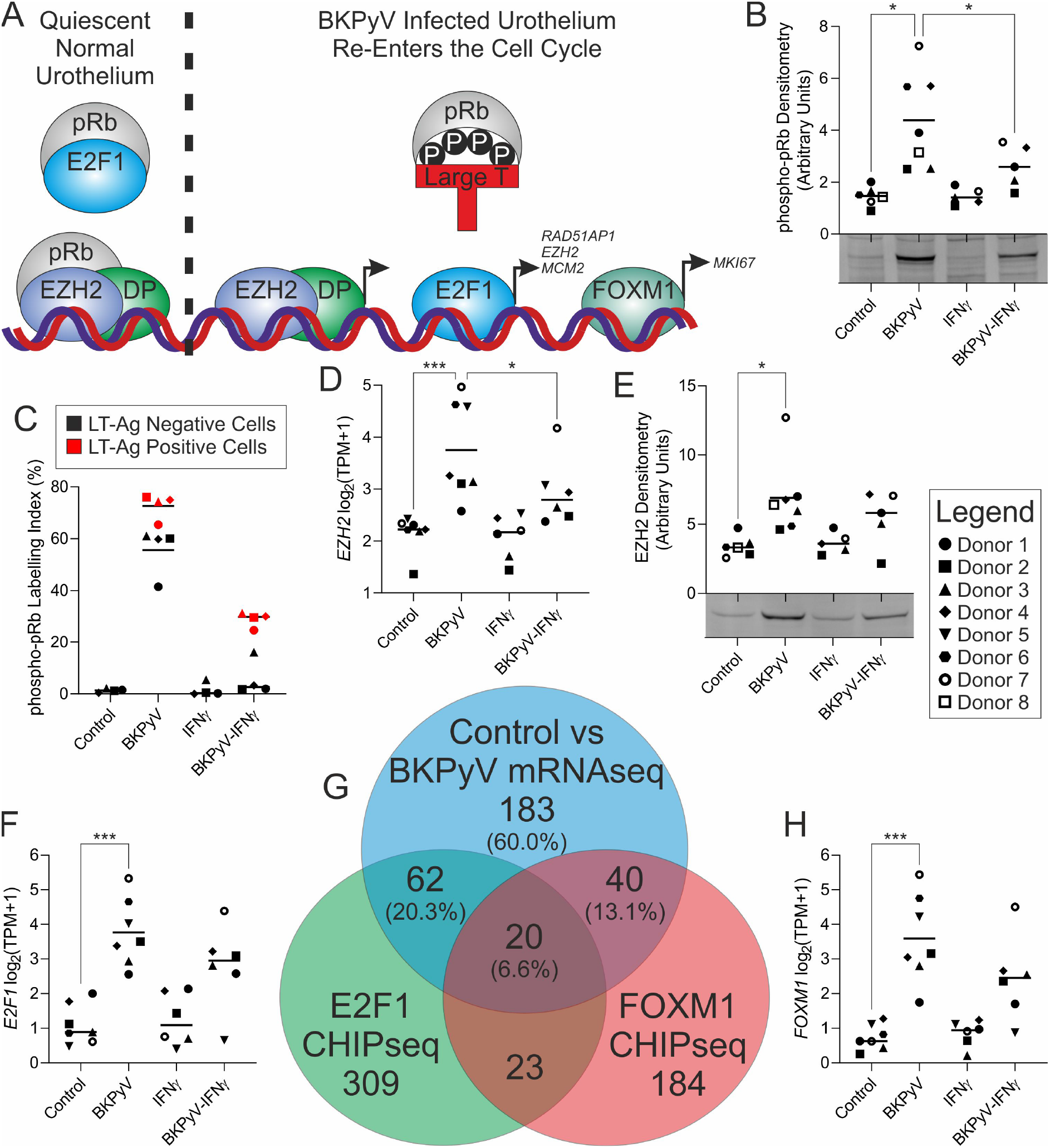
BKPyV regulates cell cycle re-entry through inactivating phosphorylation of retinoblastoma protein. (A) Schematic summary of proposed BKPyV cell cycle regulation. “pRb” = Retinoblastoma protein; “DP” = dimerization partners; “P” = phosphorylation. (B) Western blotting of retinoblastoma protein phosphorylated at serine 807/811 shows a significant infection-associated increase that was suppressed by IFNγ. (C) BKPyV infection also led to a significant increase in phosphorylated-pRb labelling index (p<0.01; Extended Data Fig. 13). When cells from infected cultures were split into LT-Ag positive and negative populations, there was mean 63.1-fold (±57.1) increase in phosphorylated-pRb labelling of LT-Ag negative cells from BKPyV-infected cultures from all donors when compared to non-infected controls. The red colouration for LT-Ag positive cells applies only to panel C. (D&E) mRNAseq and Western blotting respectively show a BKPyV-mediated increase in expression of the DREAM complex member EZH2. (F) mRNAseq shows significant induction of the retinoblastoma-regulated E2F1 transcription factor. (G) Comparison of genes significantly 2-fold induced by BKPyV infection with those reported to possess proximal E2F1^21^ and FOXM1^22^ ChIP-seq peaks, suggests 40% of the induced transcriptome may be activated by these transcription factors. (H) mRNAseq shows significant induction of the FOXM1 transcription factor by BKPyV.

The inhibition of pRb either by LT-Ag-binding or phosphorylation releases EZH2 to join the polycomb repressive complex 2 (PRC2) dimerization partners to drive transcription^20^. Involvement of EZH2/PRC2 was supported by gene-set enrichment analysis (Extended Data Fig. 12D). Transcription and protein expression of EZH2 was induced by BKPyV infection and reduced by IFNγ (Fig. 4D&E; Extended Data Fig. 14). Expression of the pRb target, *E2F1* was increased by BKPyV infection (*p*<0.001; Fig. 4F) along with other members of the E2F family (Extended Data Fig. 15). Increased E2F-activity was evidenced by gene-set enrichment of E2F1 targets in BKPyV infected cells (Extended Data Fig. 12E-G). To confirm this finding we analysed the overlap in genes associated with previously reported E2F1 ChIPseq peaks^21^ and BKPyV-induced genes and found a significant (exact hypergeometric probability *p*<4.95×10^−106^; representation factor = 38.6) overlap of 82 genes (Fig. 4G; gene lists in Extended Data Fig. 16).

Expression of the late cell cycle-associated transcription factor *FOXM1* was also induced by BKPyV infection (Fig. 4H). In agreement, GSEA supported its activity (Extended Data Fig. 12H). There was a substantial (exact hypergeometric probability *p*<1.30×10^−80^; representation factor = 43.8) overlap between genes up-regulated upon BKPyV infection and genes associated with FOXM1 binding^22^, indicative of these genes as direct targets for FOXM1 (Fig. 4G; gene lists in Extended Data Fig. 16). Taken together, increased levels and activities of E2F1 and FOXM1 accounts for 40% of the transcriptome induced by BKPyV-infection (Fig. 4G).

GSEA further revealed activation of the DNA damage response by BKPyV (Extended Data Fig. 6B) and in particular, genes implicated in homologous recombination and displacement loop structures (Extended Data Fig. 12I&J). Stabilisation of p53 protein was observed by Western blotting and indirect immunofluorescence during BKPyV infection (Fig. 5A and Extended Data Fig. 17); however, this stabilisation was not associated with increased transcription of p53 target genes consistent with LT-Ag inhibition (Fig. 5B). *RAD51* and *RAD51AP1* were significantly induced by BKPyV (mean log_2_ fold change 3.50 and 4.47, respectively; both p<0.001; Fig. 5C&D). These genes are of special interest because of their key roles in the formation of single-stranded DNA displacement loops, which could form substrates for cytosine deamination by APOBEC3 proteins. Increased Rad51 protein was confirmed by Western blotting, where a slight increase in molecular size indicated possible activating-phosphorylation of the induced Rad51 by the Chk1 kinase^23^, whose transcription was also significantly induced (Fig. 5E; Extended Data Fig. 18). Indirect immunofluorescence labelling of Rad51 identified nuclear speckles forming during BKPyV infection (Extended Data Fig. 18) and image analysis showed the appearance of nuclear speckles was significant (*p*=0.0037; Figure 5F). The LT-Ag of JCPyV was previously shown to activate the *RAD51* promoter and the two proteins were shown to co-localise in JCPyV infections^24^. Indirect immunofluorescence confirmed both the presence of nuclear Rad51 speckles in urothelial cells, and co-localisation of Rad51 with the BKPyV LT-Ag (Fig. 5F).

**Fig. 5.**
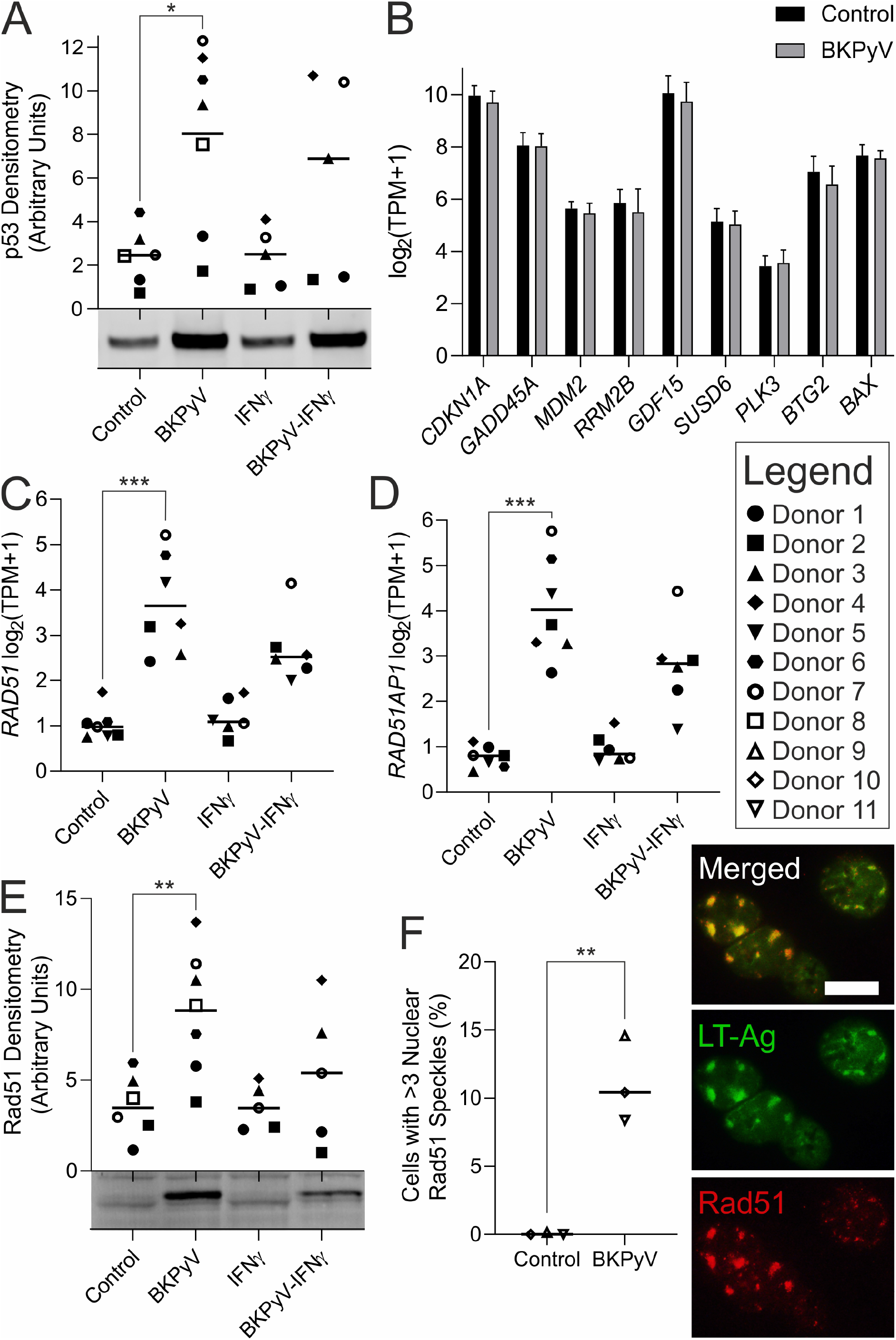
Homologous recombination was induced by BKPyV infection of human urothelium. (A) Western blotting for p53 protein showed significant stabilisation (no increase in transcription was observed) during BKPyV infection. (B) However, there was no increased expression of verified p53 target genes^25^. RAD51 and (D) RAD51AP1 transcripts were significantly induced by BKPyV infection. (E) Western blotting confirmed significant Rad51 protein induction and phosphorylation was observed as a slight increase in molecular weight when Rad51 was induced by BKPyV. (F) Analysis of indirect immunofluorescence for Rad51 found nuclear speckles were significantly increased in BKPyV infection. In some cells, indirect immunofluorescence revealed LT-Ag and Rad51 protein colocalisation to large granular deposits in the nuclei of BKPyV-infected urothelial cells. Scale bar denotes 10 μm, immunofluorescence performed on n=3 independent donors with representative images shown. Full blots and further immunofluorescence images for p53 and Rad 51 in Extended Data Fig. 17 & 18, respectively.

To explore the possible involvement of APOBEC3 cytidine deaminases (enzyme family expression data Extended Data Fig. 19), we investigated the expression of *APOBEC3A* and *APOBEC3B,* which have been implicated in the increased mutational burden in virally driven cancers, such as HPV^26^. *APOBEC3A and APOBEC3B* transcription was highly variable between donors (Fig. 6A and B). *APOBEC3A* transcript was increased by BKPyV infection in some donors (Donors 6 & 7 showed >2-fold induction) but this trend was not replicated in other donor lines (Fig. 6A). The *APOBEC3B* transcript was significantly induced >2-fold in 5 of 7 BKPyV-infected donors (mean log2 fold change 1.78 ±1.31; *p*<0.01; Fig. 6B). The *APOBEC3B* promoter was recently shown to be a direct target of pRb/E2F signalling^27^; which is consistent with the pRb/E2F and APOBEC3B induction reported here. When *APOBEC3B* expression was compared with viral transcripts, its TPM value was most significantly correlated with LT-Ag (Pearson Rho=0.98, *p*=1.407×10^−9^; Extended Data Fig. 20). However, the *APOBEC3A* transcript was not correlated with BKPyV transcripts (Extended Data Fig. 20). *APOBEC3A* and *APOBEC3B* expression can be induced by interferons, but this has been reported to be limited in urothelial cancer cell lines as compared to breast cancer lines^28^. Interestingly, IFNγ did not stimulate expression of either gene in NHU cells (Fig. 6A-D). Nor was their expression increased by IFNγ in BKPyV-infected cells (Fig. 6A-D).

**Fig. 6.**
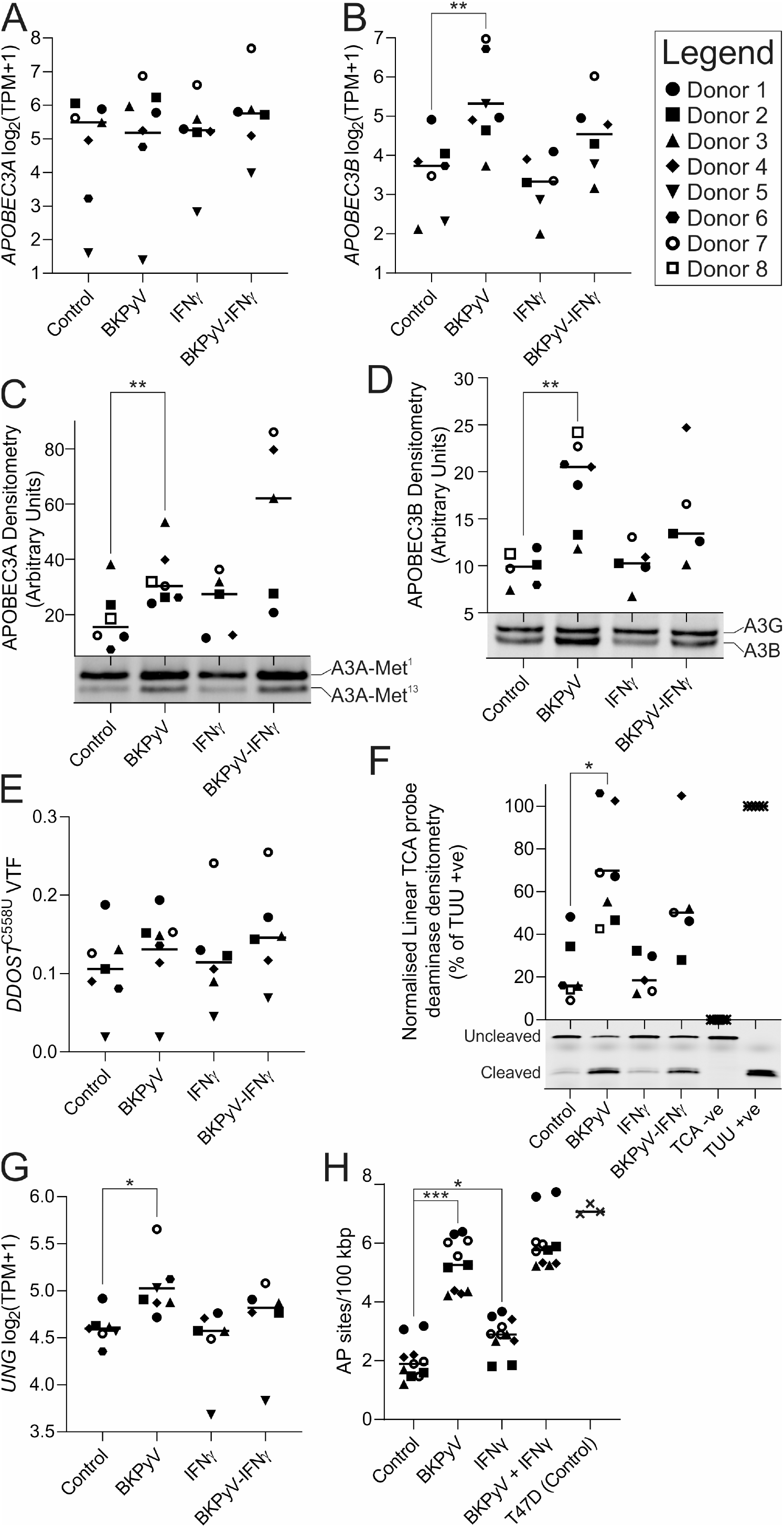
APOBEC3 expression and activity in normal human urothelium. mRNAseq expression data for APOBEC3A and APOBEC3B transcripts found only APOBEC3B was significantly (q=0.0023) induced by BKPyV infection (panel A and B, respectively). (C) Western blotting for APOBEC3A (“A3A”; full-length and truncated protein from the internal Met^13^ start site were analysed together as both possess catalytic activity ^29^) found a slight but significant (p=0.0133) induction by BKPyV infection. (D) APOBEC3B (“A3B”) and APOBEC3G (“A3G”) bands appear close together on the blots and densitometry shown here is for APOBEC3B only (as APOBE3G expression was not altered). APOBEC3B protein was significantly induced by BKPyV infection (p=0.0025). Full Western blots scans for panels C and D with densitometry for APOBEC3G are provided in Extended Data Fig. 15. (E) Variant transcript frequency (VTF) for APOBEC-mediated C>U editing of DDOST RNA at cytosine 558 quantified by mRNAseq was increased slightly, but not significantly, in 6/7 donors by BKPyV infection. Donors 5 and 7 appeared to show an IFNγ-mediated induction of APOBEC3A-function. (F) Deaminase assays for NHU protein lysates against a linear RTCA probe preferentially targeted by APOBEC3B confirm showed significant induction of activity following BKPyV infection. Significance was tested using a Wilcoxon matched-pairs signed rank test (p=0.0313). Full deaminase gel scans are provided in Extended Data Fig. 17. Deaminase assays were also performed against a YTCA hairpin probe, in the presence and absence of RNA, to evaluate APOBEC3A activity (Extended Data Fig. 18). (G) mRNAseq expression data for the uracil-DNA glycosylase gene UNG found it was significantly induced by BKPyV infection (q=0.0213). (H) Apurinic/apyrimidinic (AP) sites assay. The assay was performed on DNA from five independent donors with 2-3 independent cultures per donor. T74D breast cancer cells were included as a positive control known to generate apurinic/apyrimidinic sites in their genomes ^32^. The legend in the top right identifies the donor by specific point shapes in all dot plot panels.

APOBEC3 protein abundance was studied by Western blotting (Fig. 6C&D, Extended Data Fig. 21). APOBEC3A has two enzymatically-active isoforms, with a smaller variant generated by internal translation initiation at a methionine at position 13 (Met^13^)^29^. Both APOBEC3A isoforms were increased in all donors following BKPyV infection (*p*=0.0025; Fig. 6C), although there was large variation in the magnitude of increase. In cells from Donors 4 & 5, APOBEC3A protein abundance in BKPyV-infected cultures was increased >2-fold by IFNγ exposure (Fig. 6C). APOBEC3B was significantly increased by BKPyV infection (*p*=0.0054; Fig. 6D). Unexpectedly, IFNγ slightly reduced APOBEC3B protein abundance in BKPYV-infected cultures (Fig. 6D). Western blot densitometry was significantly correlated with TPM for both APOBEC3A and APOBEC3B (*p*=0.0147 and *p*=0.0003, respectively). Densitometry analysis suggested that during BKPyV infection, the ratio of APOBEC3A to APOBEC3B protein was 1.95:1 (±1.18; n=7).

A recent study identified RNA-editing of the *DDOST* transcript at cytosine 558 to be a common activity of APOBEC3A that was not observed with APOBEC3B^30^. *DDOST* transcript editing was mildly increased in 6/7 donors by BKPyV infection (25.2% increase ±23.9; Fig. 6E). Across all mRNAseq samples (n=26) *DDOST* C558U variant transcript frequency (VTF) was significantly correlated with *APOBEC3A* TPM (Pearson Rho=0.803; *p*=5.205×10^−7^) but not *APOBEC3B* TPM (Pearson Rho=0.278; *p*=0.167; Extended Data Fig. 22).

APOBEC3-activity was assessed by deaminase assays using single stranded DNA probes conjugated to fluorochromes (Fig. 6F). BKPyV infection significantly increased deaminase activity against a linear probe with an RTCA motif that is the substrate preference of APOBEC3B (mean log_2_ fold change 1.66 ±1.05; *p*=0.0303; Fig. 6F, Extended Data Fig. 23). Linear regression analysis confirmed significant relationships between the deaminase activity and *APOBEC3B* protein abundance (F=33.19, df=21; *p*<0.0001) and to a lesser extent *APOBEC3A* protein (F=5.030, df=21; *p*=0.0358) (Extended Data Fig. 24). Further studies using a hairpin probe assay, designed to favour APOBEC3A-activity^31^ also showed increased activity following BKPyV-infection. The addition of RNA, demonstrated to inhibit APOBEC3B, but not APOBEC3A activity^31^, had no effect on deamination of the hairpin substrate, suggesting that APOBEC3A-activity was induced by BKPyV infection (Extended Data Fig. 25).

The nature of APOBEC mutagenesis, being widely distributed throughout the genome in mostly extra-genic regions, combined with the lack of clonal expansion in this experimental model meant that it was not possible to confirm APOBEC-activity through an RNA-derived mutational signature (Extended Data Fig. 26). APOBEC-activity in the genome converts cytosine to uracil, which requires excision by the uracil DNA glycosylase enzyme that leaves an apurinic/apyrimidinic (AP) site suitable for subsequent repair. The uracil-DNA glycosylase enzyme gene “*UNG*” was significantly induced by BKPyV-infection (p<0.05; Fig. 6G). Furthermore, the increased deaminase activity observed in BKPyV infection (Fig. 6F) was associated with significant (*p*<0.001) damage to the host genome, measured as an increase in AP sites (Fig. 6H). Linear regression analysis confirmed a significant relationship between AP sites and APOBEC3B protein abundance (F=9.033, df=17; *p=*0.008) but not APOBEC3A (F=1.190, df=17; *p*=0.185; Extended Data Fig. 27).

To understand how APOBEC3-activity is relates to viral infection, proximity ligation assays were used to study protein:protein interactions. LT-Ag is a critical biological effector of the BKPyV life-cycle and has previously been shown to interact directly with pRb^33^. Here we used the pRb:LT-Ag interaction as a positive control for proximity ligation assays and included ZO3:LT-Ag as a negative control (zonula occludins 3 (ZO3) is a differentiated urothelial tight junction member and not known to enter the nucleus; Fig. 7). LT-Ag appeared to co-localise with Rad51 by indirect immunofluorescence (Fig. 5F) and proximity ligation assays confirmed that the two proteins were frequently within <40 nm of one another (Fig. 7). In addition, APOBEC3 enzymes co-localised with LT-Ag in the nuclei of infected urothelial cells (Fig. 7). LT-Ag has been shown to bind dsDNA and act as a helicase; however, the precise nature of LT-Ag’s binding partners and complex membership remains to be resolved. It is tempting to hypothesise that LT-Ag acts as a lynchpin bringing multiple factors together around displacement loops to harness DNA repair enzymes for viral genome replication but additionally leading to collateral damage of the host genome (summarised Fig. 8).

**Fig. 7.**
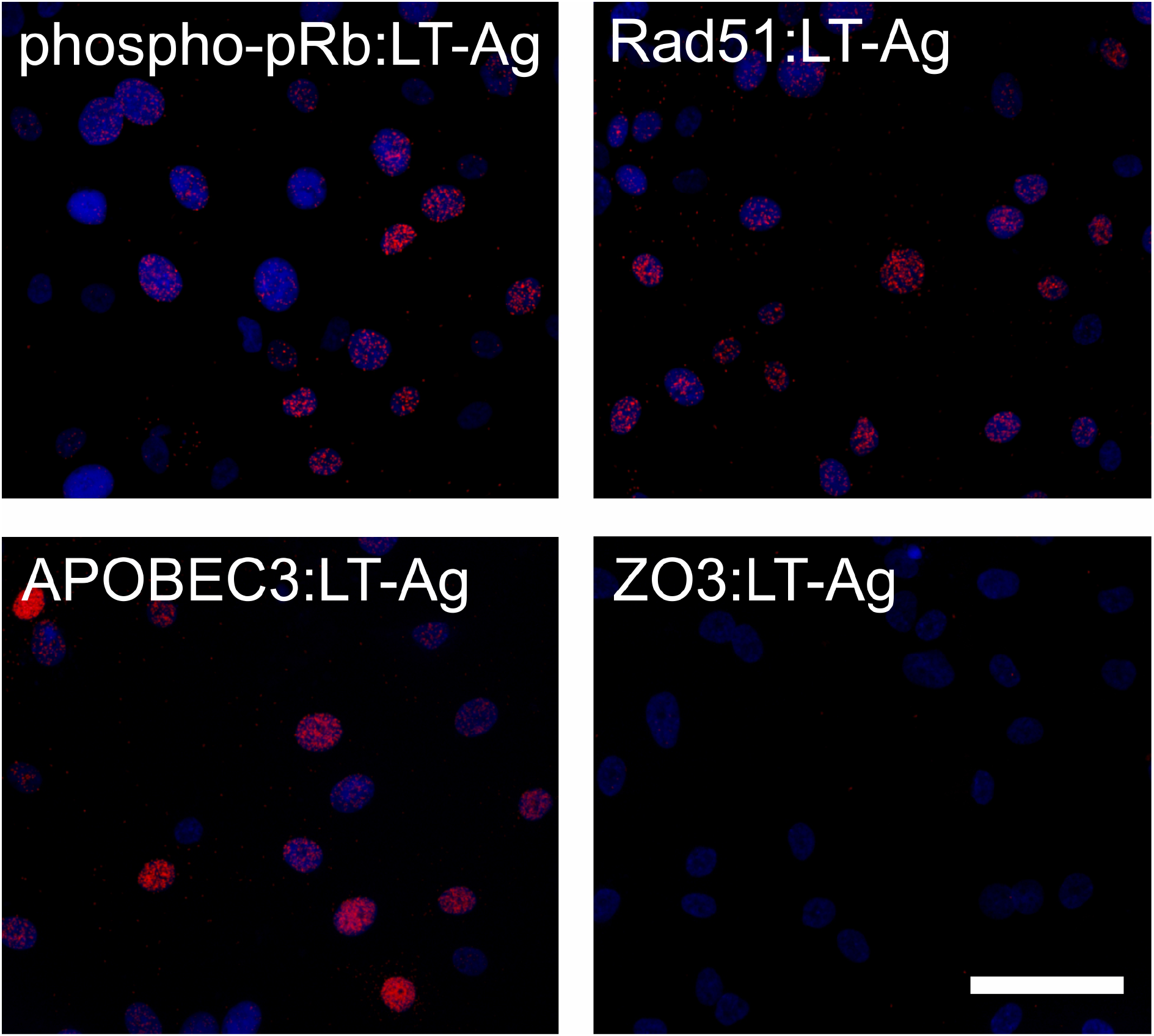
Proximity ligation assays suggested that during BKPyV infection of human urothelium, nuclear complexes form involving large T antigen (LT-Ag) and phospho-Retinoblastoma (phopho-pRb), Rad51 and the APOBEC3 proteins. Zonula occludins 3 (ZO3) is a tight junction protein critical for differentiated urothelial barrier formation and expressed in this tissue model at cell:cell junctions included here as a negative control. Scale bar denotes 50 μm, n=3 donors (Extended Data Fig.28-30).

**Fig. 8.**
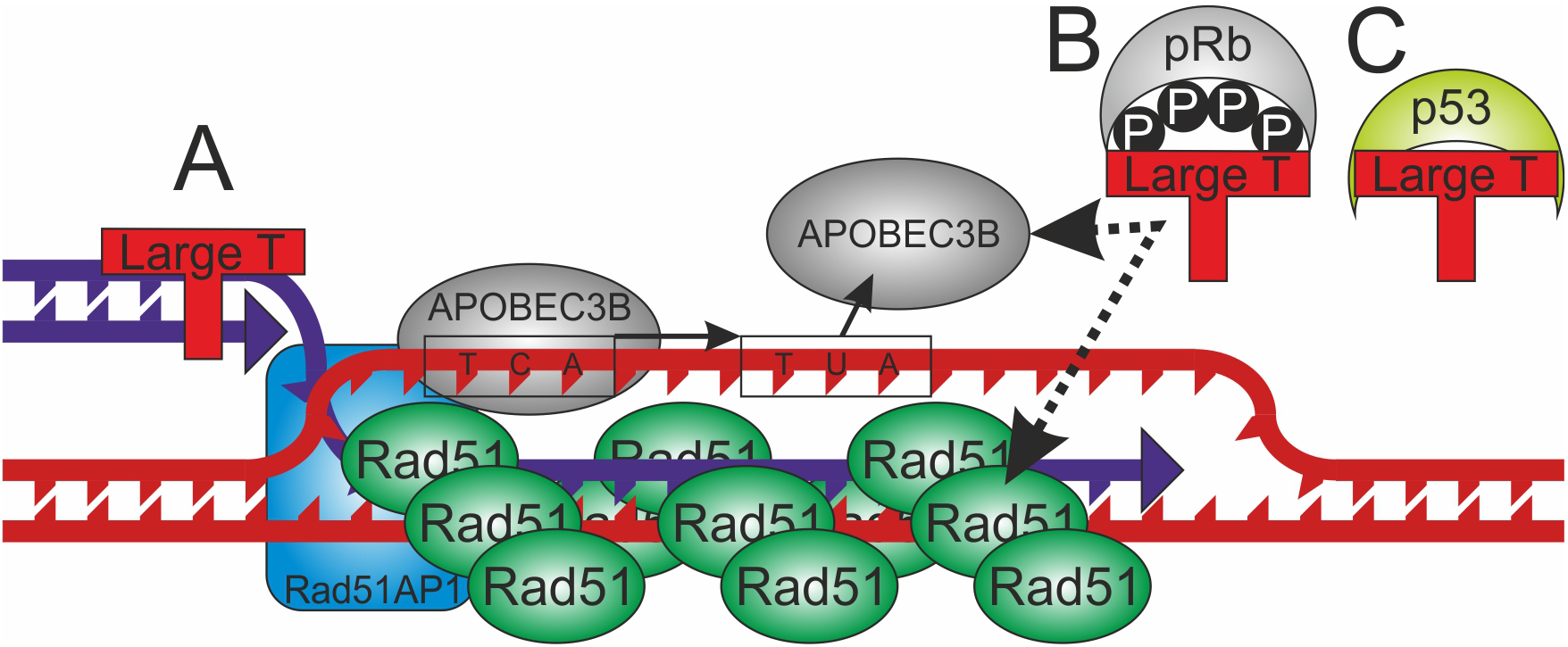
Schematic of hypothetical model of urothelial DNA damage at displacement loops at the G2 checkpoint during BKPyV infection. LT-Ag interactions might derive from (A) its DNA-binding helicase activity or (B) one of the other dimerization domains present in the LT-Ag protein. (C) LT-Ag inhibition of p53 is likely important for this process as p53 would normally prevent Rad51 oligomerisation by direct binding^41^. The evidence from this study is most robust for BKPyV-induced APOBEC3B activity but APOBEC3A protein was consistently more abundant during infection and its role requires further validation.

## Discussion

Tracking the molecular events leading to human epithelial malignancy can be challenging when the carcinogenic event may have occurred decades in the past. Genomic mutational signatures offer insight but, in the case of bladder cancer, controversy exists between the putative epidemiological risk from smoking and mutational signatures found in tumours^1, 7^, which suggest a missing viral agency. Using a tissue-mimetic *in vitro* model that replicates both the barrier and mitotic quiescence of human urothelium *in situ*, we here provide the first experimental evidence that BKPyV can directly infect differentiated human urothelium, driving the host cells out of mitotic quiescence to facilitate the viral life cycle. This is achieved by viral LT-Ag mediated inactivation of the host tumour suppression machinery, including p53 and pRB. A key corollary of these events is the acquisition of host genomic damage through APOBEC3A/B activity, which we associate with DNA displacement loops formed during homologous recombination at the G2/M checkpoint (Summarised Fig. 8). This evidence establishes BKPyV as an infectious agent with the capacity to damage the urothelial genome with the potential to initiate carcinogenesis.

Indirect evidence supporting the relevance of these pathways comes from observations that BLCAs are more common in immune-supressed patients following solid organ transplant (standardised incidence ratio 1.52^34^) and in particular kidney transplant (meta-analysis standardised incidence ratio 2.46^35^). Indeed, where a renal transplant patient tests positive for BKPyV activity, one retrospective study of male transplant patients found the risk of BLCA increased >8-fold^36^. The time from renal transplant to BLCA diagnosis was an average of 9.6 years in a recent study^37^, comparable with the known disease course in HPV-driven tumours. Other biological processes leading to immune-insufficiency, including ageing, may also be sufficient to trigger BKPyV reactivation and lead to transient reinfection of the urothelium, driving host cell-cycle activity and genomic damage. In light of this, we propose due consideration be given to BKPyV vaccine development^38^ and subsequent trials with an initial focus on renal transplant patients as an at-risk population where significant benefits in terms of graft outcomes and urinary tract pathologies (including cancer) are likely. Viewed through the prism of BKPyV as a risk factor for BLCA, a number of features of the disease that are not currently understood (including risk factor interactions^39^ and pronounced gender differences^40^) might start to be explained.

## Methods

### Normal Human Urothelial (NHU) Cell Culture

Eleven independent NHU cell lines of finite (non-immortalised) lifespan were used in this study. The cell lines were established as described^42^ using anonymous discarded tissue from renal transplant surgery, with NHS Research Ethics Committee approval. NHU cells were propagated in Keratinocyte Serum-Free Medium (KSFM; 0.09mM Ca^2+^) supplemented with bovine pituitary extract, recombinant human EGF and cholera toxin. Following expansion, NHU cells were differentiated in medium supplemented with adult bovine serum and [Ca^2+^] elevated to 2mM, according to published methods^15^.

### BKPyV Infection and IFNγ Treatment

BKPyV Dunlop Strain was expanded for use in renal proximal tubule epithelial cell cultures which were scrape harvested at 14 days post infection, sonicated and frozen as aliquots at −80 °C. Multiplicity of infection (MOI) was calculated by fluorescent focus unit assay using IncuCyte ZOOM analysis (Essen BioScience, Ann Arbor, MI, USA), as previously reported^43^. All cell cultures in this study were infected with 0.45 μm filtered BKPyV containing medium at MOI=1 for 3-4 hours at 37 °C before virus containing medium was removed and cultures continued.

At 7 days post infection (dpi) some cultures were exposed to IFNγ (200U/mL, BioTechne #285-IF) for a further seven days to mimic an immune response to the infection.

### mRNA Analysis

Total RNA was collected in TRIzol reagent (Invitrogen). The relative abundance of selected transcripts was assessed by Reverse Transcribed - quantitative Polymerase Chain Reaction (RT-qPCR) using the following forward and reverse primers (all 5’-3’) to amplify *LT-Ag* (GAGTAGCTCAGAGGTGCCAACC and CATCACTGGCAAACATATCTTCATGGC^44^), *VP1* (CTTTGCTGTAGGTGGAGAACCC and CTCCTGTGAAAGTCCCAAAATAC^45^) and *GAPDH* (CAAGGTCATCCATGACAACTTTG and GGGCCATCCACAGTCTTCTG). Primers were optimised to give a linear response over a 1,000 fold dilution range and a single product as characterised by a single peak in the dissociation curve. Amplification was monitored using SYBR Green dye on a QuantStudio™ 3 Real-Time PCR System machine (ThermoFisher). All measurements were performed in triplicate and calculated using the ΔΔct method relative to GAPDH expression.

Samples from 14dpi were selected for mRNA sequencing (mRNAseq) using the Illumina NovaSeq 6000 generating 150bp paired-end reads (Novogene UK, Cambridge, UK). All mRNAseq data has been deposited at GSEXXX. Following standard quality control, gene-level expression values in transcripts per million (TPM) were derived against the Gencode v35 human transcriptome using kallisto v0.46.1^46^. For analysis of the BKPyV transcriptome (Fig. 2A), sequences (derived from BKPyV reference; GenBank NC_001538.1) were appended to the human transcriptome to generate “relative TPMs” as a measure of viral transcript abundance. Reads were also aligned to the human (GRCh38) and BKPyV reference genome assemblies with HISAT2 v2.2.0^47^, and single nucleotide variants detected following best practices using GATK v4.1.0^48^, PicardTools v2.20.0^49^, SAMtools v1.10^50^ and VCFtools v0.1.15^51^. Mutational signatures were then processed as previously described^1^.

Differentially expressed genes were identified using the sleuth v0.30.0^52^ implementation of the likelihood ratio test (LRT), accounting for matched genetic backgrounds, generating Benjamini-Hochberg corrected q-values. For volcano plots (performed in R v4.0.4 EnhancedVolcano v1.8.0), fold change values used a TPM+1 transformation to reduce the influence of low abundance transcripts. To compare manipulations of the culture model, π-values were calculated as described elsewhere^53^:

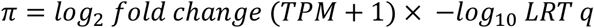

Gene set enrichment analysis (GSEA) was performed using π-values (derived from the Sleuth LRT q-values) and the pre-ranked list feature implemented in the python package GSEApy (0.10.2; available at: https://github.com/zqfang/GSEApy). The ranked list of genes was run against the four following Molecular Signatures Database (MSigDB) collections: hallmarks (h.all.v.7.2.symbols.gmt), curated (c2.all.v.7.2.symbols.gmt), gene ontology (c5.all.v.7.2.symbols.gmt), and oncogenic signatures (c6.all.v.7.2.symbols.gmt) (available at: https://data.broadinstitute.org/gsea-msigdb/msigdb/release/7.2/).” Overlap between genes significantly 2-fold induced by BKPyV-infection and previously reported E2F1/FOXM1 ChIPseq peaks^21, 22^ was assessed by calculating the exact hypergeometric probability.

### Indirect Immunofluoresence

NHU cell cultures on glass 12-well slides were fixed in methanol:acetone (50:50 v/v) 30 seconds, air dried and stored frozen.

Primary antibodies, applied overnight at 4°C, were anti-SV40 LT-Ag (1:200, mouse monoclonal “Pab 108”, Santa Cruz Biotechnology, sc-148), anti-VP1 (1:100, mouse “pab597”, kind gift from Chris Buck at the National Institute of Health, Bethesda), anti-Rad51 (1:1,000, rabbit “EPR4030(3)”, Abcam #ab133534), anti-Ki67 (1:400, mouse, “MM1”, Leica, NCL-L-Ki67-MM1), anti-MCM2 (1:500, rabbit, “D7G11”, Cell Signalling #3619), anti-Phospho-pRb Serine 608/807/811 (1:500, rabbit, Cell Signalling #9308), anti-p53 (1:40, mouse, “D01”, Cell Signalling #18032), anti-ZO3 (1:800, rabbit, Cell Signalling #3704) and anti-APOBEC3A/B/G (1:100 rabbit monoclonal, clone 5210-87-13^54^).

Unbound primary antibodies were removed by washing in phosphate-buffer saline (PBS) and secondary antibodies (Goat-anti-Mouse Alexa-488 and Goat-anti-Rabbit Alexa-594, Molecular Probes) were applied for 1 h at ambient temperature. Slides were washed in PBS, with 0.1 μg/ml Hoechst 33258 added to the penultimate wash, before mounting in ProLong Gold Antifade Mountant (ThermoFisher) and visualisation by epifluorescence on a BX60 microscope (Olympus).

Image analysis was performed in ImageJ (v1.53c Java 1.8.0_172) by creating regions of interest (ROI) around the nuclei in images of Hoechst 33258 DNA staining. Corresponding images of antibody labelling were subsequently overlaid using the ROI manager. To derive labelling indices, nuclear intensity in a minimum of 1,000 cells was calculated and a threshold for labelling intensity was established on an appropriate negative control sample. Analysis of Rad51 nuclear speckles was performed using the Speckle Inspector tool from the BioVoxxel Toolbox (v2.5.1) in a minimum of 1,000 cells. Nuclei with three speckles or fewer were disregarded.

### Proximity Ligation Assays (PLA)

NHU cell cultures on glass 12-well slides were fixed in 10% neutral buffered formalin for 10 minutes before permeabilisation in PBS (phosphate-buffered saline) containing 0.5% (w/v) Triton X-100 for 30 minutes. PLA were performed using the Duolink^®^ In Situ Red kit as per the manufacturer’s instructions (Sigma). Finally, slides were fixed in Methanol:Acetone (50:50 v/v) for 30 seconds and air-dried before mounting in Duolink^®^ In Situ Mounting Medium with DAPI. Epifluorescence was visualised on a BX60 microscope (Olympus).

### Protein Lysate Collection

Cell cultures in 10cm dishes were scrape-harvested in 500 μL lysis buffer containing 0.2% (v/v) protease inhibitors (Protease Inhibitor Cocktail set III, Calbiochem). Lysis buffer comprised: 25 mM HEPES-KOH (pH 7.5), 10% glycerol, 150 mM NaCl, 0.5% Triton X-100, 1 mM ethylenediaminetetraacetic acid. Lysates were sonicated and clarified by centrifugation at 13,000 g. A bicinchoninic acid (BCA) protein assay (ThermoScientific) was used to normalise loading into western blots and deaminase assays.

### Western Blotting

Fifty μg of protein lysate per lane was resolved on NuPAGE gels using the Novex electrophoresis system (Invitrogen) at 200 V. Electrotransfer to PVDF-FL membranes (Millipore) was completed in a Tris–glycine buffer at 30 V for 2 h at 4°C before appropriate blocking. The test antibodies used were anti-SV40 LT-Ag (1:250, mouse monoclonal “Pab 108”, Santa Cruz Biotechnology, sc-148), anti-VP1 (1:250, mouse “pab597”, kind gift from Chris Buck at the National Institute of Health, Bethesda), anti-MCM2 (1:1,000, rabbit, “D7G11”, Cell Signalling #3619), anti-Phospho-pRb Serine 608/807/811 (1:1,000, rabbit, Cell Signalling #9308), anti-EZH2 (1:1,000, rabbit “D2C9”, Cell Signalling #5246), anti-p53 (1:1,000, mouse, “D01”, Cell Signalling #18032), anti-Rad51 (1:10,000, rabbit “EPR4030(3)”, Abcam #ab133534), anti-APOBEC3A/B/G (1:800 rabbit monoclonal, clone 5210-87-13^54^). Homogeneous loading and transfer were evaluated using β-actin antibodies (Sigma, Clone AC15, Mouse, 1:10,000 dilution). Membranes were labelled with the appropriate IRDye conjugated secondary antibody (LI-COR) at ambient temperature for 1 h and visualised by epifluorescent infrared illumination at 700 and/or 800 nm using the Odyssey Sa scanner and software (LI-COR). Densitometry was performed using Image Studio Lite Ver 5.0 software (LI-COR). Cropped Western blots are shown in the main Fig.s with full blots provided as Extended Data Fig.s 4-6. Western blots were loaded with equal protein amount in every lane and correct loading/transfer was confirmed by probing for β–actin (Extended Data Fig. 4).

### Deaminase Activity Assays

Protein lysate [1μg/μL] was RNase A digested at 37°C for 15 min following addition of 1 μg RNase A (Qiagen) per 25 μg protein. In Extended Data Fig. 19, the RNase digestion was omitted and 100 ng/μL of urothelial RNA was added to achieve greater selectivity for APOBEC3A (as previously described ^31^). 10 μg protein lysate was mixed with 1 pmol ssDNA substrate (IDT, Germany) and 0.75 U uracil-DNA glycosylase (UDG; New England Biolabs) in a total volume of 12 μL and incubated at 37 °C for 1 h. 10 μl 1M NaOH was added and samples incubated for 15 min at 37 °C. Finally, 10 μl 1M HCl was added to neutralize the reaction and samples were separated by electrophoresis through 15% urea-polyacrylamide gel electrophoresis gels in Tris-borate-EDTA (1×) at 150V for 2 h. Gels were visualised by epifluorescent infrared illumination at 700 nm using the Odyssey Sa scanner and software (LI-COR). Densitometry was performed using Image Studio Lite Ver 5.0 software (LI-COR). A positive TUU probe and negative control (TCA probe without lysate) were included to aid experimental interpretation.

The ssDNA substrates used in these assays were:

1. Linear RTCA /5IRDye700/A*T*A*ATAATAATAATAATAATAAT**<u>ATCA</u>**ATAATAATAATAATAATA*A*T*A
2. Linear TUU positive control probe /5IRDye700/A*T*A*ATAATAATAATAATAATAAT**<u>ATUU</u>**ATAATAATAATAATAATA*A*T*A
3. Hairpin YTCA (as previously described “oTM-814”^31^ to be selective for APOBEC3A-mediated deamination in the presence of exogenous RNA) /5IRD700/TTTTATTTTGCAATTG**<u>TTCA</u>**ATTGCAAAATTT*G*T*T

Asterisks in the DNA probes denote phosphorothioate modifications, which confer resistance to both endo- and exonucleases, providing increased oligo stability. A graphical description of the method is provided as Extended Data Fig. 31.

### Apurinic/apyrimidinic Sites Assay

Genomic DNA was extracted from 2-3 independent cultures of NHU cells from each of 5 independent donors using Nucleospin Tissue (Machery-Nagel) spin columns. Apurinic/apyrimidinic (AP) sites were quantified using the Oxiselect DNA damage ELISA kit (AP sites; STA-324; Cell Biolabs Inc. San Diego, CA, USA), according to manufacturer’s instructions. A standard curve of aldehyde reactive probe DNA was used to quantify the number of genomic AP sites. The assay was performed using T47D breast cancer cell DNA as a positive control for AP sites, as we have previously described^32^.

### Statistical Analysis

Data were assessed for statistical significance using Prism 8.3.0 software (Graphpad). On all graphs statistical *p* or *q* value significance is represented as follows; * <0.05, ** <0.01 & *** <0.001.

## Data Availability Statement

All mRNAseq data that support the findings of this study have been deposited in the NCBI GEO database with the accession code GSEXXX (to be released upon publication).

## Code Availability Statement

No unique code was used in this manuscript and all code employed is freely available from the cited sources.

## Acknowledgements

This study was funded by York Against Cancer.

## Notes

### Competing Interest Statement

The authors have declared no competing interest.

