## Extended Data - Figures for "BK polyomavirus (BKPyV) is a risk factor for bladder cancer through induction of APOBEC3-mediated genomic damage"

**Supplementary Information**

Established bladder cancer cell lines sourced from ATCC/ECACC were tested and free of *Mycoplasma spp.* and were authenticated by short tandem repeat profiling using the PowerPlex16 System (Promega) within 5 passages of use in this study; all cell lines were a perfect match to the ATCC/ECACC genotype records. Cell lines were all cultured for this study in DMEM:RPMI 1640 (50:50, v:v) with 5% fetal bovine serum.

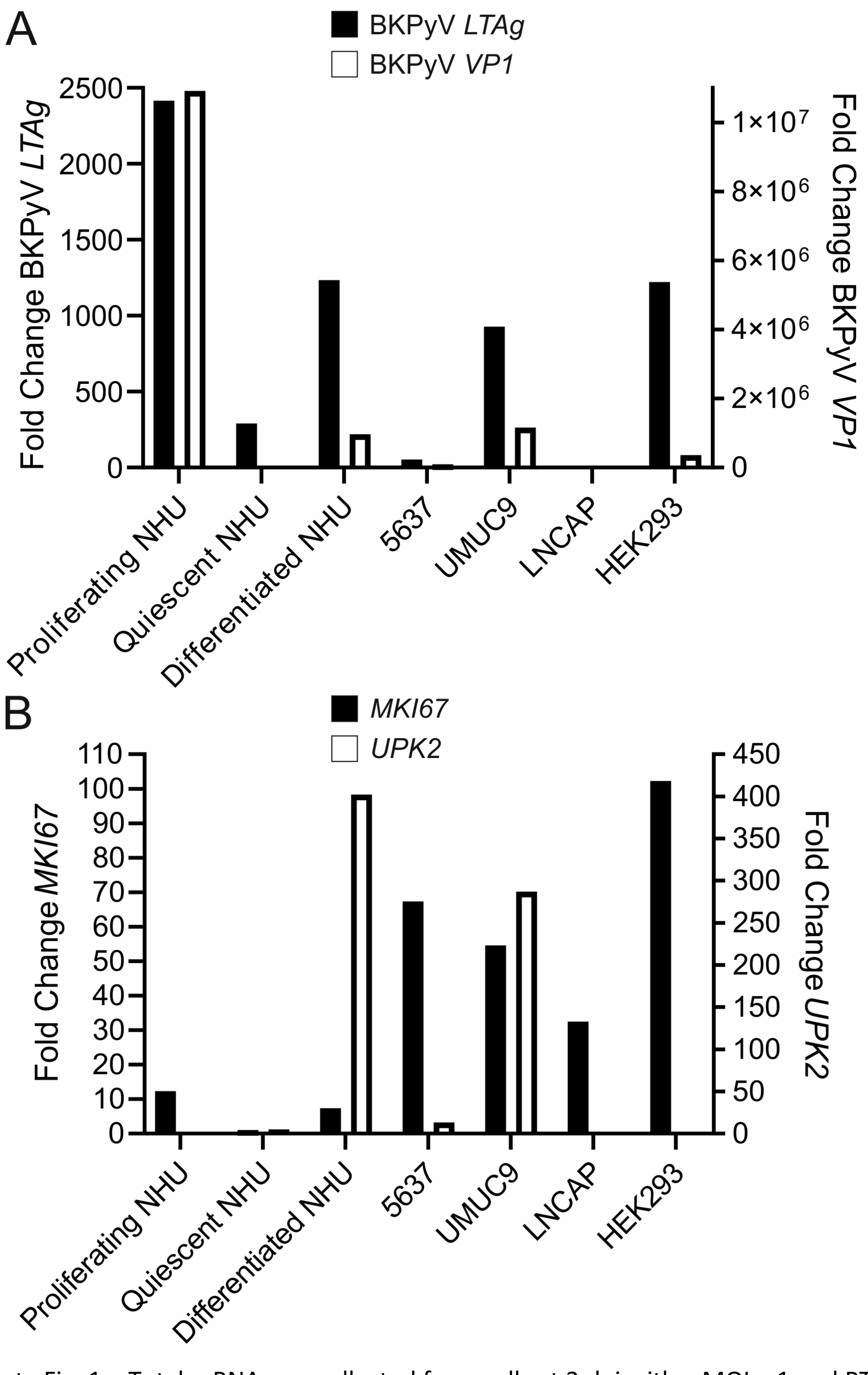


Extended Data Fig. 1 – Total mRNA was collected from cells at 3 dpi with a MOI = 1 and RT-qPCR was performed for the early (*LT-Ag*) and late (*VP1*) BKPyV transcripts (Panel A); and *MKI67* and *UPK2* host transcripts (Panel B). BKPyV preferentially infected differentiated urothelium (indicated by uroplakin 2/*UPK2* expression; panel B) whether normal or neoplastic. BKPyV readily infected undifferentiated proliferating NHU cells where the viral life cycle proceeded to late promoter driven *VP1* transcript expression at 3dpi. Differentiated NHU cells are mitotically quiescent and were therefore compared with undifferentiated "quiescent NHU" cells, where quiescence was achieved by contact inhibition. 5637 and UMUC9 were respectively chosen as poorly-differentiated and well-differentiated models of urothelial carcinoma (based on *UPK2* expression). LNCAP and HEK293 cells were included as negative and positive control models for BKPyV infection, respectively.

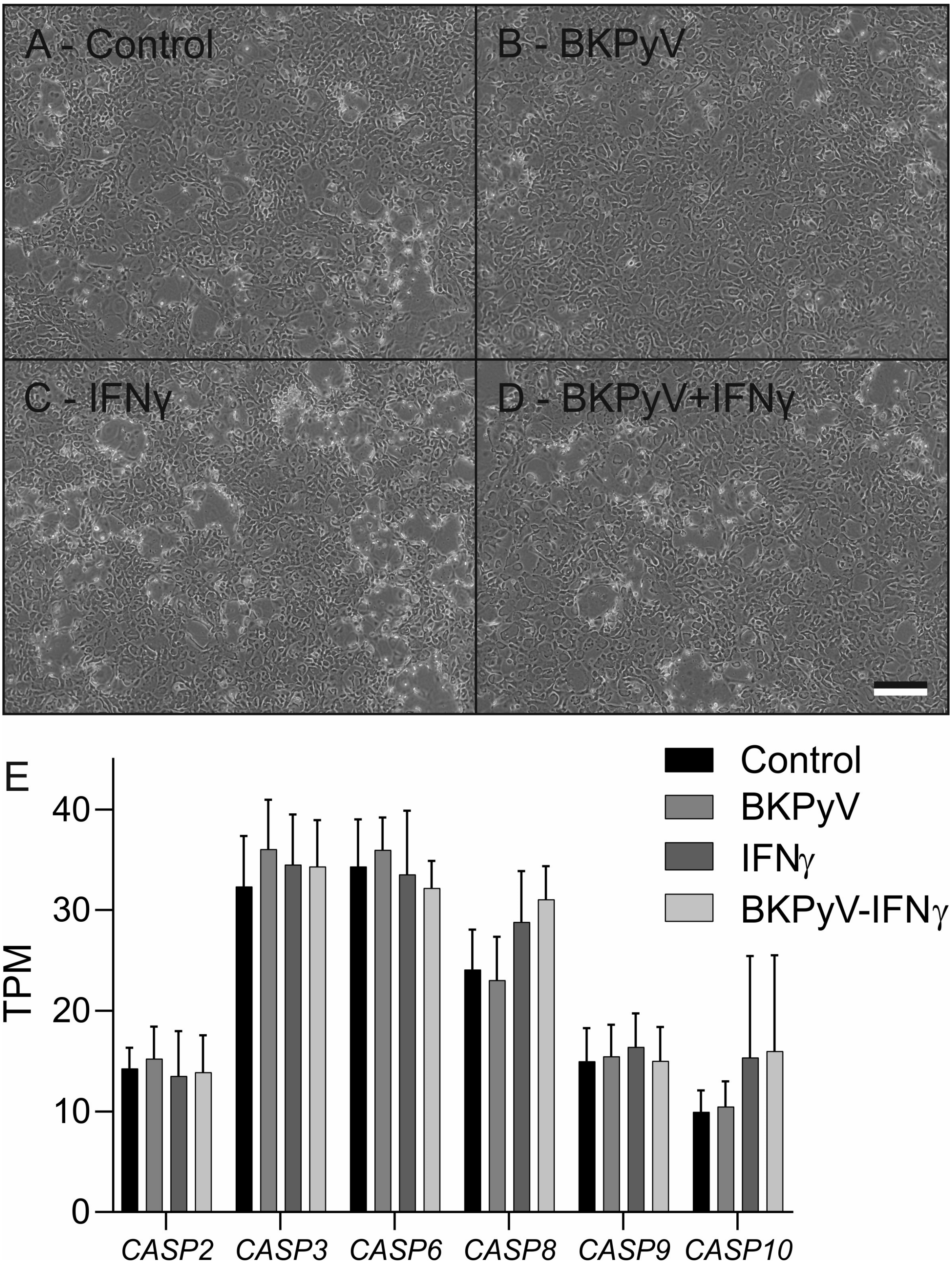


Extended Data Fig. 2 – (A-D) Example phase contrast images of one donor line of the studied differentiated normal human urothelial (NHU) cells at 14dpi showing no signs of apoptosis or cell loss in the cultures. The cultures are 100% confluent but stratification of the urothelium into multi-layered tissues takes them out of the plane of focus in some areas, giving the appearance of bare plastic. Scale bar in panel D denotes 200μm. (E) Transcriptomic analysis of the pro-apoptotic caspases found the maximum mean increase in expression was 12% (*CASP3*; n=6/7).

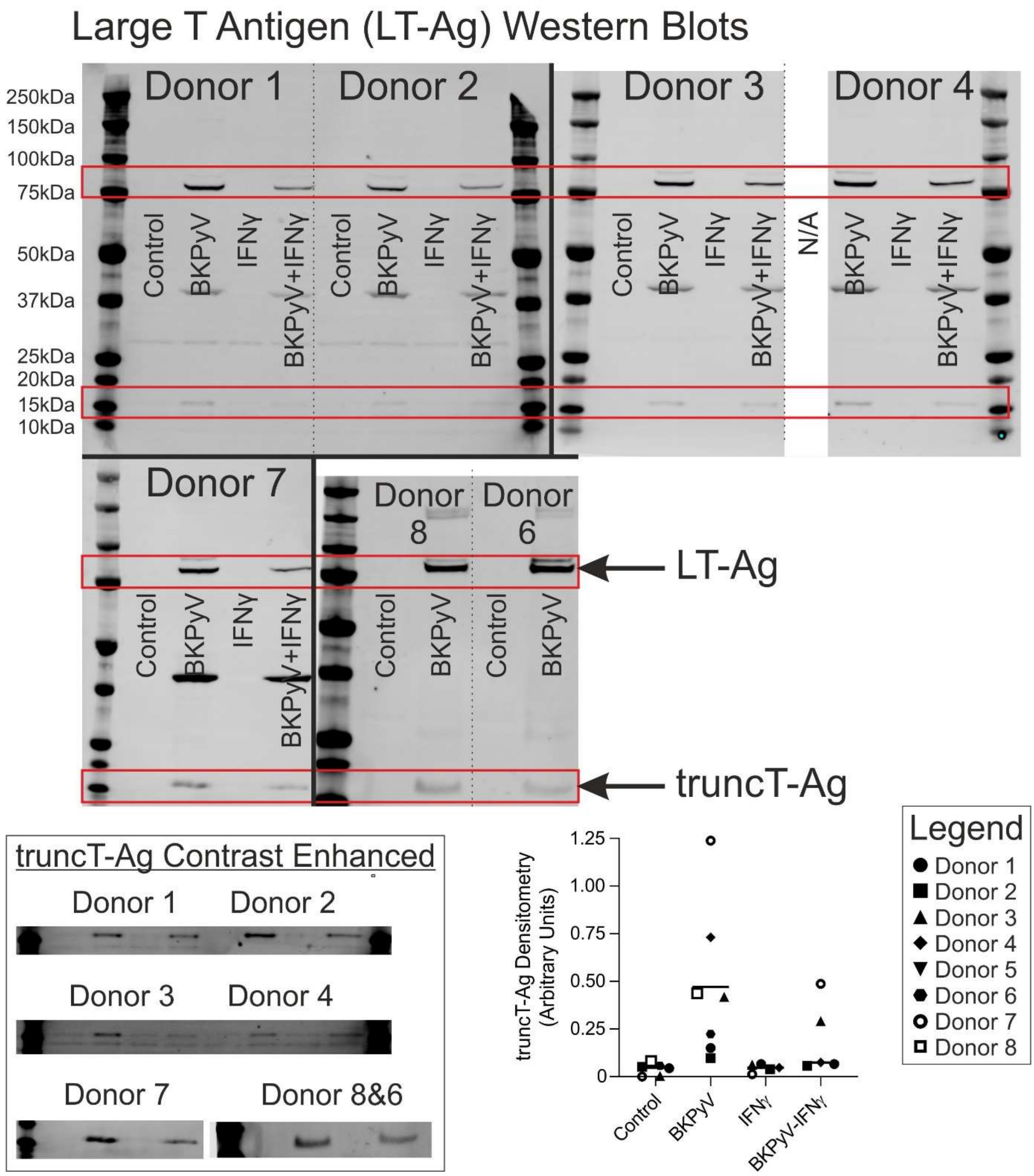


Extended Data Fig. 3 – Full anti-LT-Ag Western blots used for densitometry in Fig. 2. Predicted molecular weight for BKPyV large T antigen is 80.5kDa (UniProtKB - P03071). Lower 40kDa band visible on some blots is retained VP1 from previous probing of the Western blot shown in Extended Data Fig. 5. The band at 17-20 kDa is the truncated of T-antigen[1] which is expressed to a lesser extent than full length LT-Ag. The box shows contrast enhancement of the truncT-Ag band on all blots and the dot plot shows densitometry for the truncT-Ag band. Like LT-Ag, truncT-Ag expression was reduced by IFNγ. The control cells for Donor 4 were lost to an infection during culture and were therefore not available (N/A) for analysis.

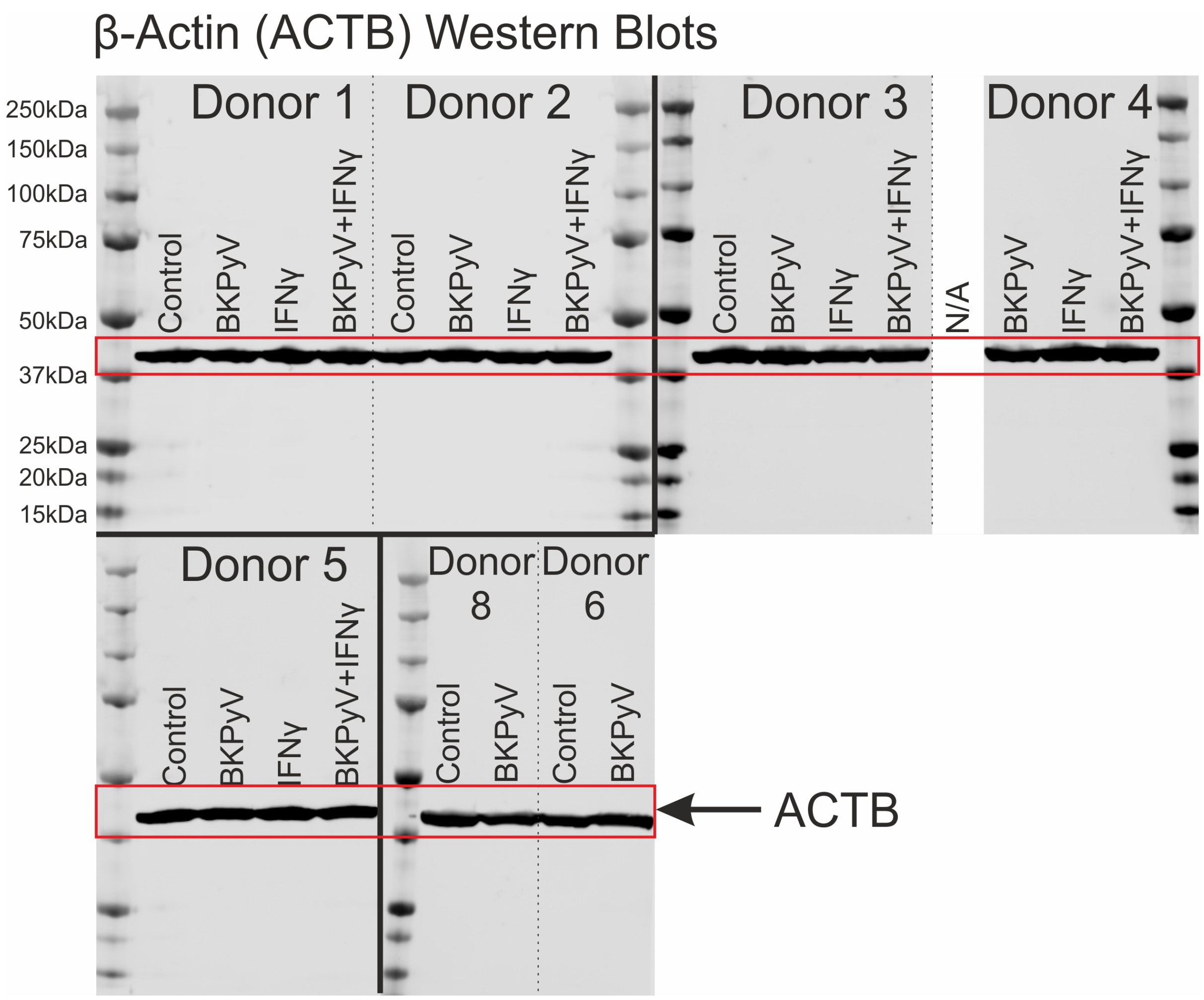


Extended Data Fig. 4 – The loading of Western blots was normalised by running the same amount of protein per lane (50µg). Here we show, Western blotting of β–actin (as a housekeeping protein not expected to change in abundance) as additional evidence, supporting even loading and transfer of protein using this method. The control cells for Donor 4 were lost to an infection during culture and were therefore not available (N/A) for analysis.

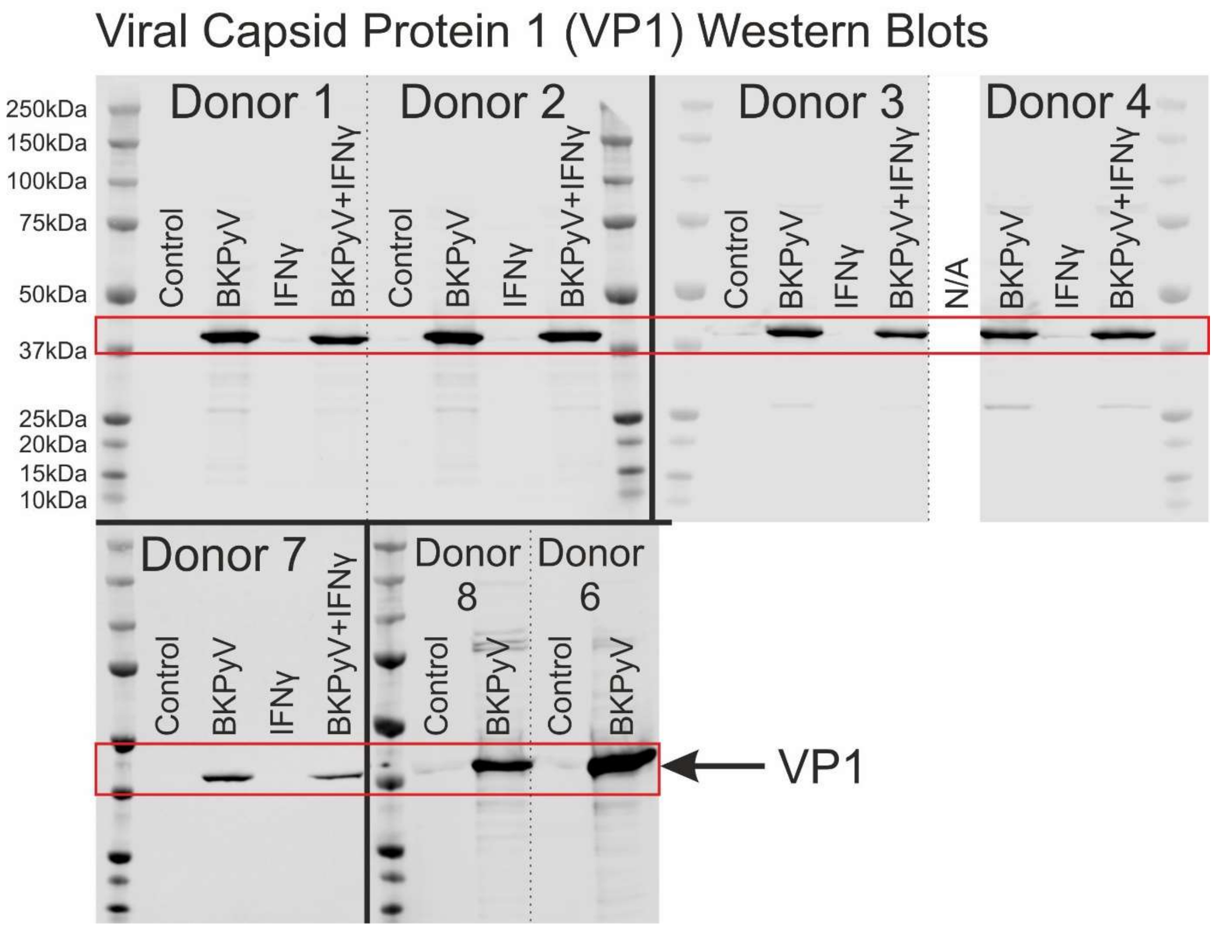


Extended Data Fig. 5 – Full anti-VP1 Western blots used for densitometry in Fig. 2. Predicted molecular weight for BKPyV VP1 capsid protein is 40.1kDa (UniProtKB - P03088). The control cells for Donor 4 were lost to an infection during culture and were therefore not available (N/A) for analysis.

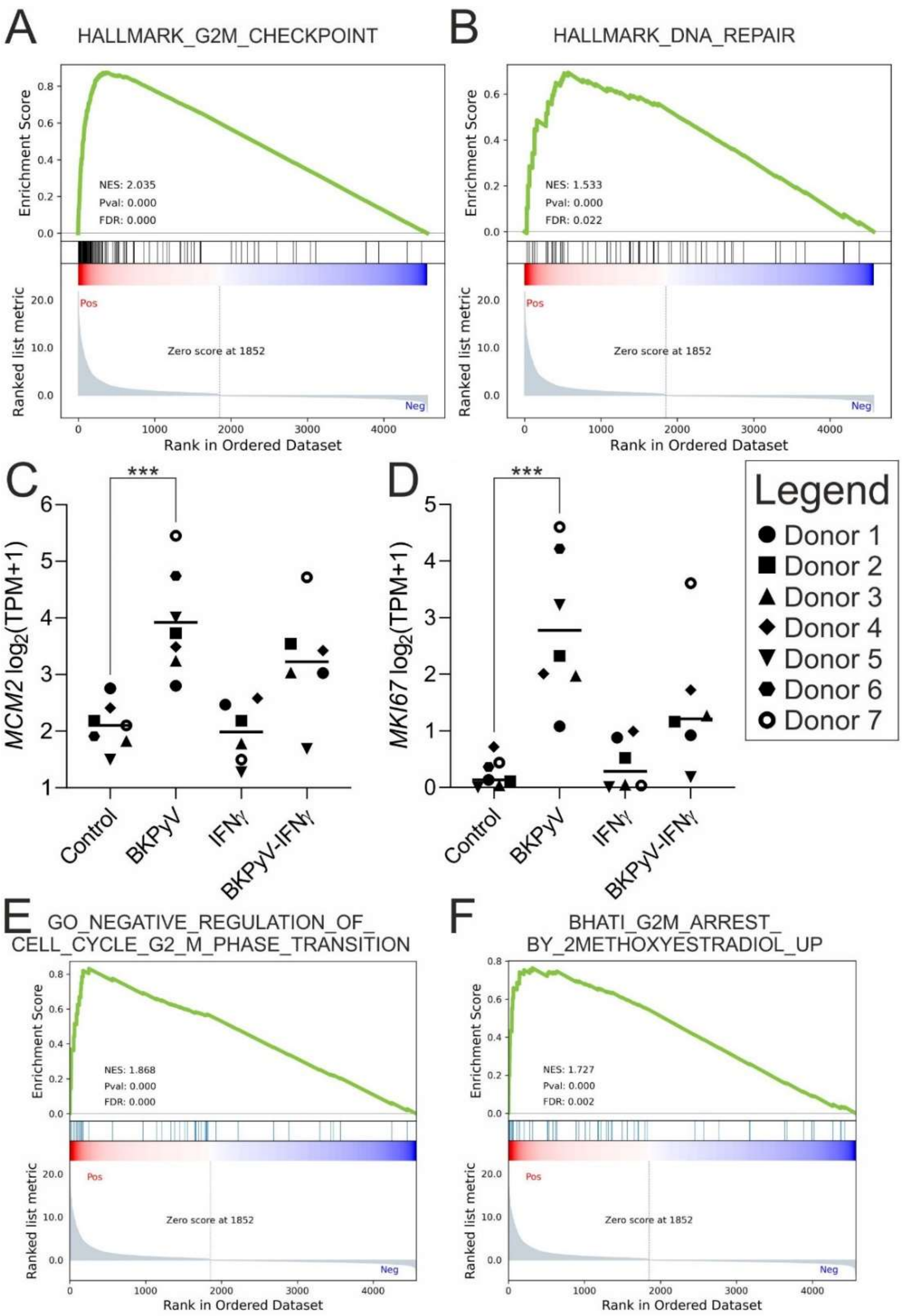


Extended Data Fig. 6 – Gene-set enrichment analysis (GSEA) of mRNAseq π-values for the control vs BKPyV comparison revealed increased expression related to the G2/M checkpoint of the cell cycle (A) and DNA repair processes (B). mRNAseq data for the proliferation markers (C) MCM2 (D) MKI67. MCM2 transcript expression is not completely lost in quiescent cultures as was observed at the protein level (Fig. 3). MKI67 is traditionally used as a marker for cells in active cell cycle and is lost as cells exit the cycle into G0. Both MCM2 and MKI67 transcript expression was significantly ($p<0.001$) induced by BKPyV infection (C&D). Interferon-γ exposure reduced MCM2 and MKI67 transcript expression in BKPyV-infected cultures from all donors tested compared with infection alone; however, the variance in reduction made this change not statistically significant (C&D). GSEA of mRNAseq π-values for the control vs BKPyV comparison also revealed increased expression associated with negative regulation of the G2 to M transition (E) and experimentally-induced G2-arrest by 2-methoxyestradiol (F).

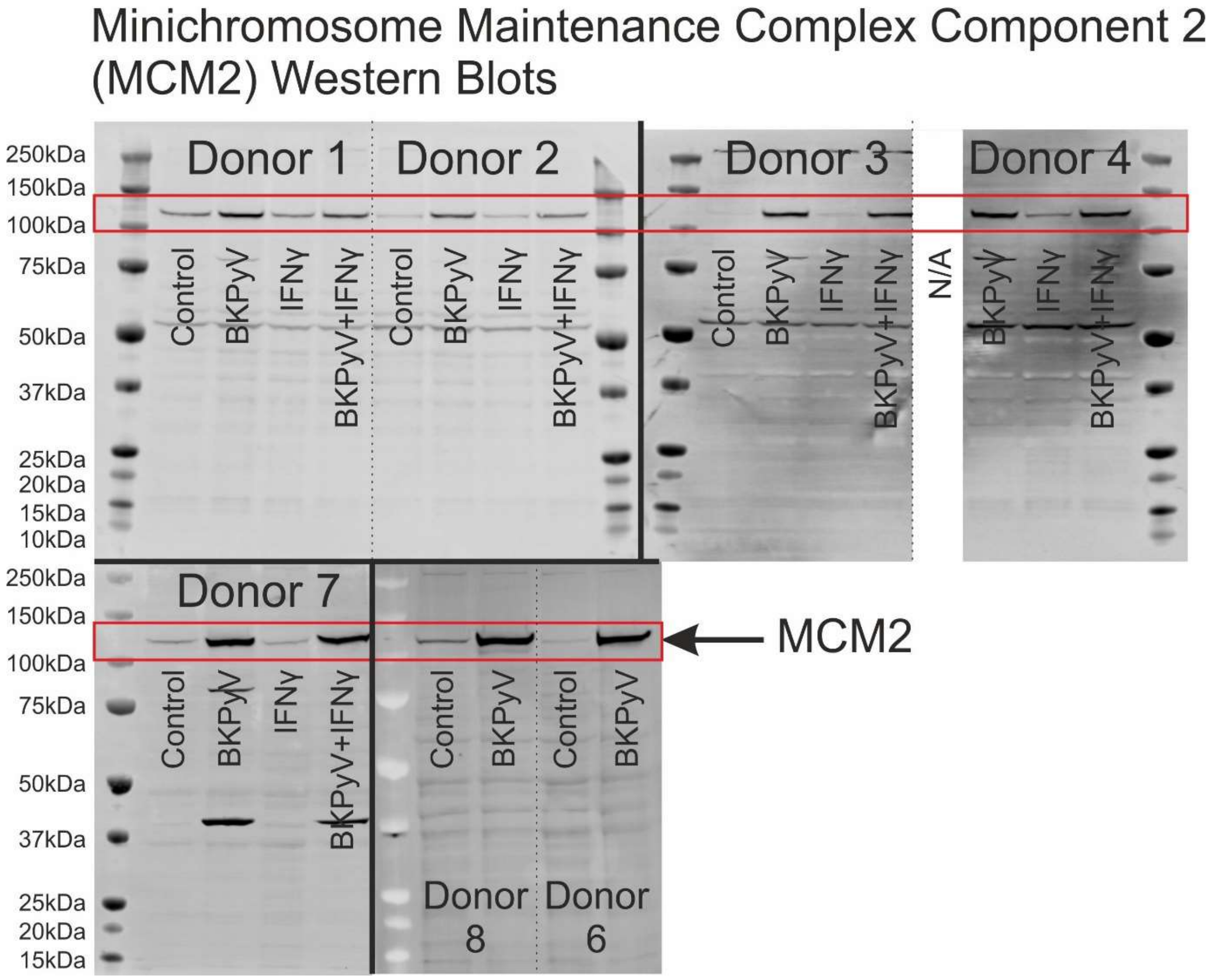


Extended Data Fig. 7 – Full anti-MCM2 Western blots used for densitometry in Fig. 3. Predicted molecular weight for DNA replication licensing factor MCM2 is 101.9kDa (UniProtKB - P49736). The lower ~55kDa band (Donor 1-4) was considered non-specific and the 40kDa band on the Donor 5 blot is residual antibody from prior probing for VP1. The control cells for Donor 4 were lost to an infection during culture and were therefore not available (N/A) for analysis.

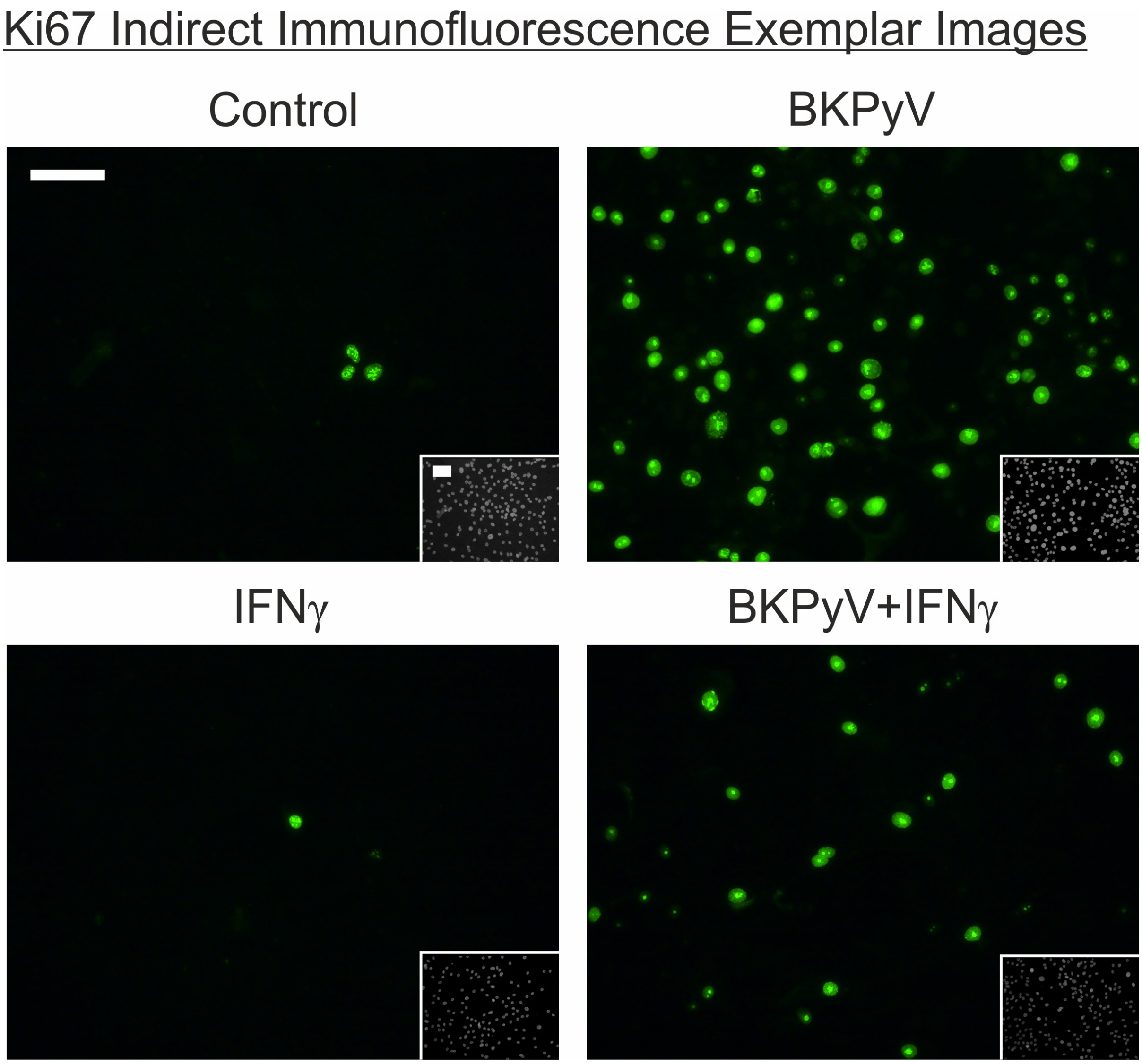


Extended Data Fig. 8 – Indirect immunofluorescence labelling of Ki67 in NHU cell cultures. Ki67 positive nuclei in the control cultures show the many small speckles characteristic of the G1 cell cycle stage. By contrast, Ki67 positive nuclei in the BKPyV infected cultures display the few larger nucleolar granules of labelling that identifies the G2 stage of the cell cycle. Inset shows nuclei stained with Hoechst 33258. White scale bar in control main panel and inset denotes 100 μm.

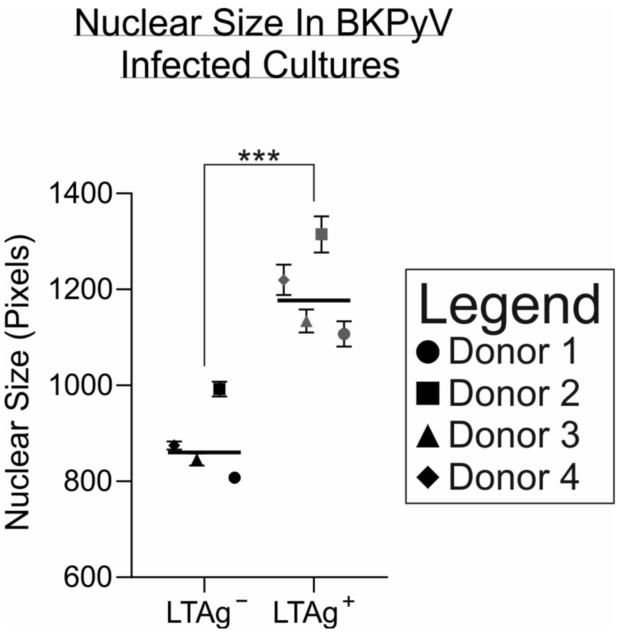


Extended Data Fig. 9 – Hoechst 33258 staining of urothelial cells in BKPyV infected cultures, showed a significant ($p$=0.0001) increase in the size of nuclei for LT-Ag labelling positive cells as measured by pixel area (line indicates the mean nuclear size in n=4 independent donors with >1,000 cells analysed in each condition).

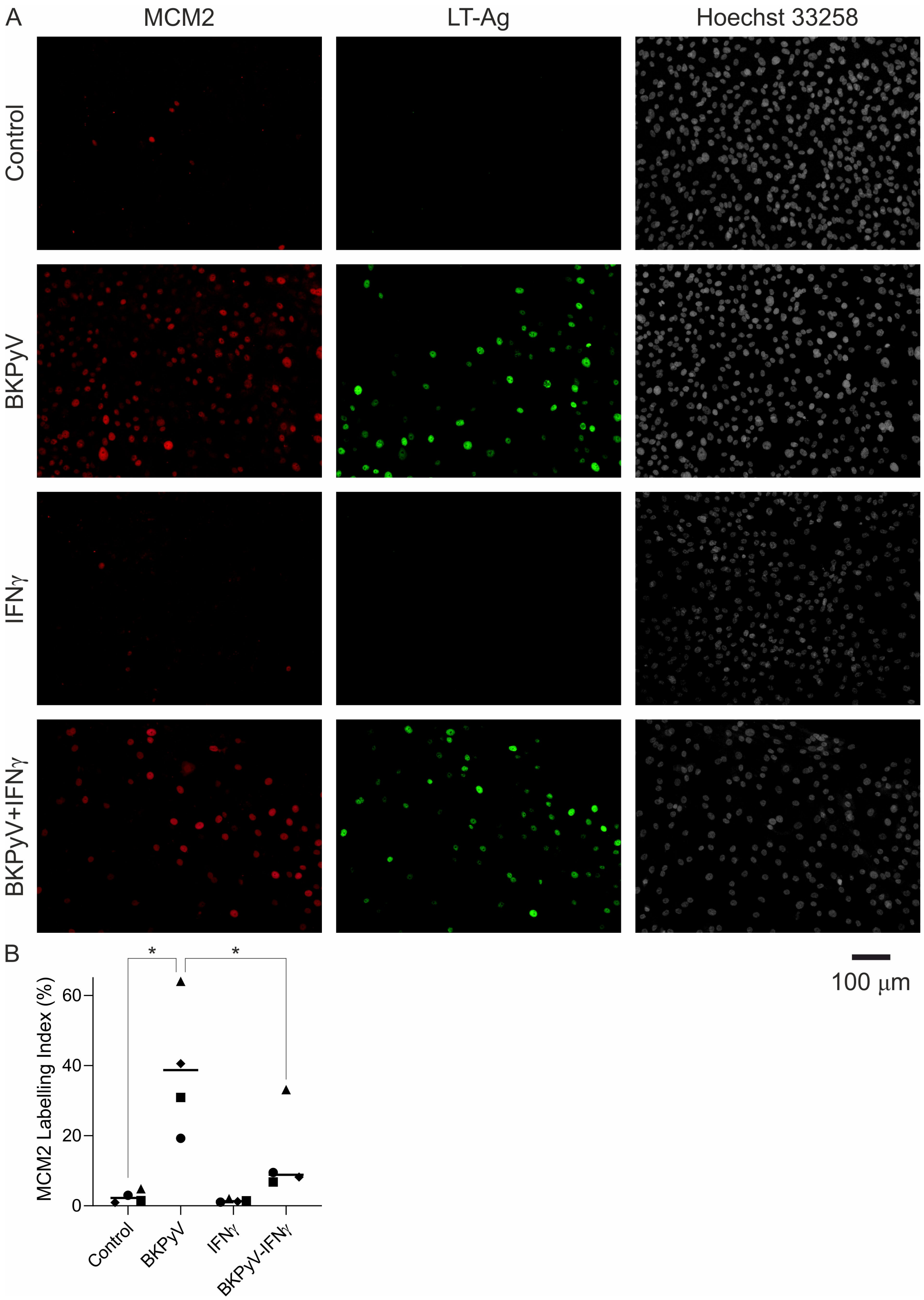


Extended Data Fig. 10 – (A) Indirect Immunofluorescence labelling for MCM2 (red) is shown alongside LT-Ag (green) labelling of the same cells. Hoescht 33258 DNA staining is included in grayscale. In BKPyV-infected cells there was MCM2 positivity in the absence of LT-Ag labelling. (B) MCM2 labelling indices showed a significant increase in positive cells in BKPyV infected cultures. In addition, the number of MCM2 positive cells in BKPyV-infected cultures was significantly lower when IFNγ was added to the culture (line indicates the mean nuclear size in n=4 independent donors with >1,000 cells analysed in each condition).

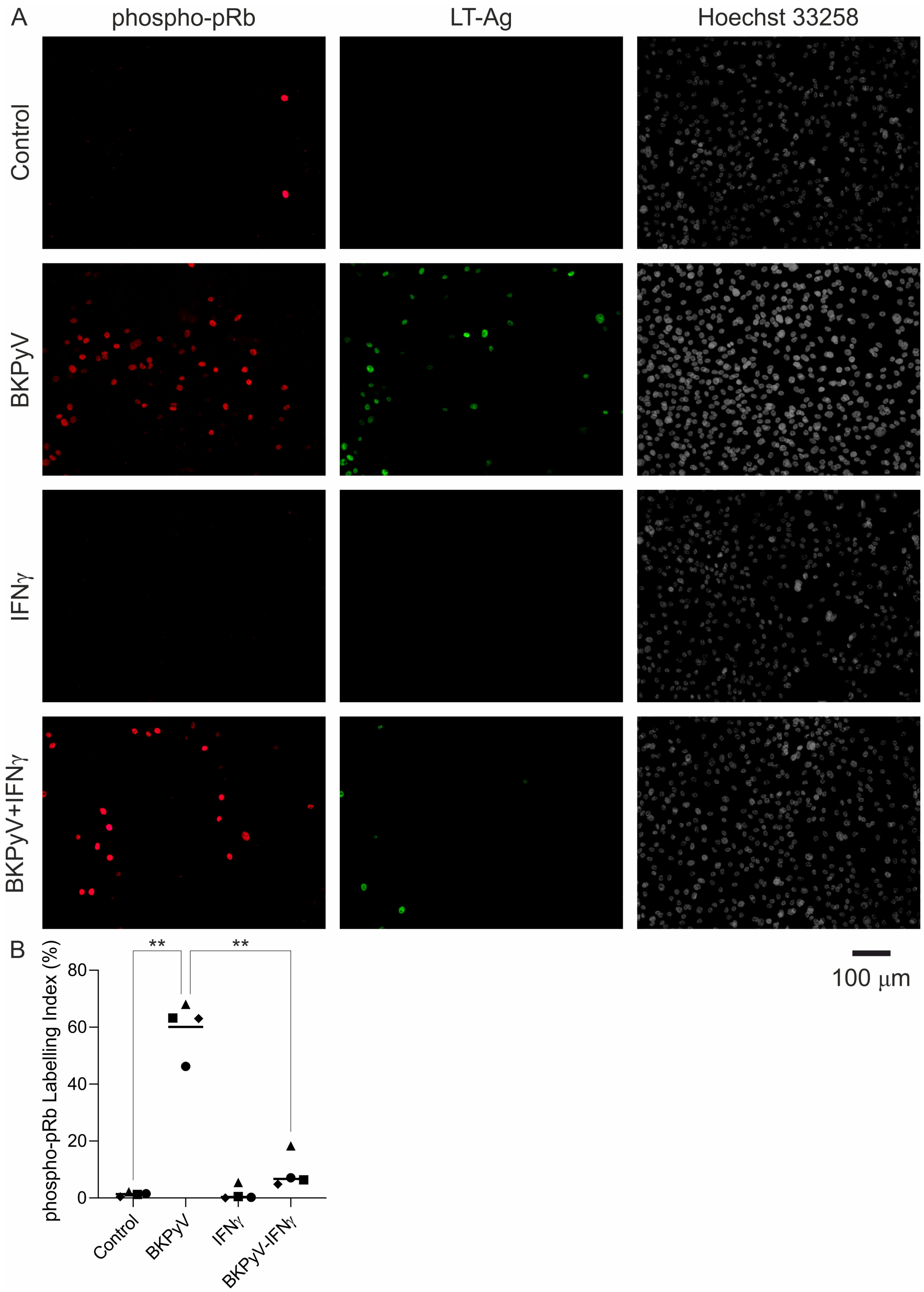


Extended Data Fig. 11 – (A) Indirect Immunofluorescence labelling for phosphorylated-Retinoblastoma (phospho-pRb; red) is shown alongside LT-Ag (green) labelling of the same cells. In BKPyV-infected cells there was frequent phospho-pRb positivity in the absence of LT-Ag labelling. (B) phospho-pRb labelling indices showed a significant increase in positive cells in BKPyV infected cultures. In addition, the number of phospho-pRb positive cells in BKPyV-infected cultures was significantly lower when IFNγ was added to the culture (line indicates the mean nuclear size in n=4 independent donors with >1,000 cells analysed in each condition).

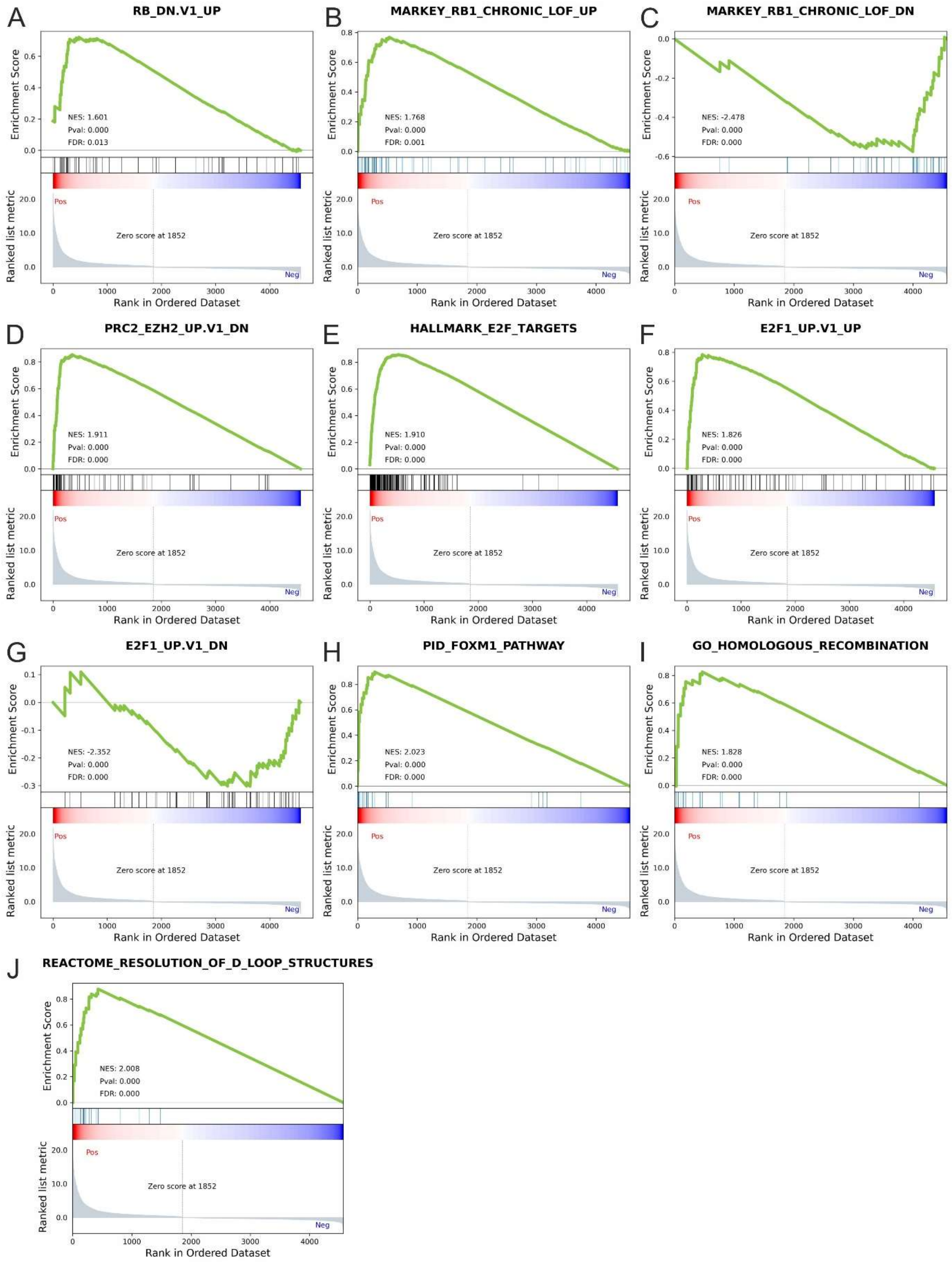


Extended Data Fig. 12 – Gene-set enrichment analysis (GSEA) of mRNAseq π-values for the control vs BKPyV comparison.

(A) RB_DN.V1_UP = gene set derived from epidermal-specific ablation of Rb gene.

(B) MARKEY_RB1_CHRONIC_LOF_UP = Genes up-regulated in mouse embryonic fibroblasts isolated from RB1 knockout mice: chronic loss of function (LOF) of RB1

(C) MARKEY_RB1_CHRONIC_LOF_DN = Genes down-regulated in mouse embryonic fibroblasts isolated from RB1 knockout mice leading to chronic loss of function (LOF) of RB1

(D) PRC2_EZH2_UP.V1_DN = Genes down-regulated in TIG3 cells (fibroblasts) upon knockdown of EZH2 gene. BKPyV infection increased EZH2 transcript and protein (Fig. 4D&E). Thus genes that went down when EZH2 was knocked down in fibroblasts, also go up when EZH2 expression goes up in BKPyV infection.

(E) HALLMARK_E2F_TARGETS = Genes encoding cell cycle related targets of E2F transcription factors.

(F) E2F1_UP.V1_UP = Genes up-regulated in mouse fibroblasts over-expressing E2F1

(G) E2F1_UP.V1_DN = Genes down-regulated in mouse fibroblasts over-expressing E2F1

(H) PID_FOXM1_PATHWAY = Pathway Interaction Database (NCI, NIH and Nature Publishing Group) FOXM1 transcription factor network

(I) GO_HOMOLOGOUS_RECOMBINATION = Gene Ontology Consortium contributed gene set associated with a DNA recombination process that results in the equal exchange of genetic material between the recombining DNA molecules.

(J) REACTOME_RESOLUTION_OF_D_LOOP_STRUCTURES = Reactome contributed list of genes associated with resolution of D-loop Structures through Synthesis-Dependent Strand Annealing (SDSA).

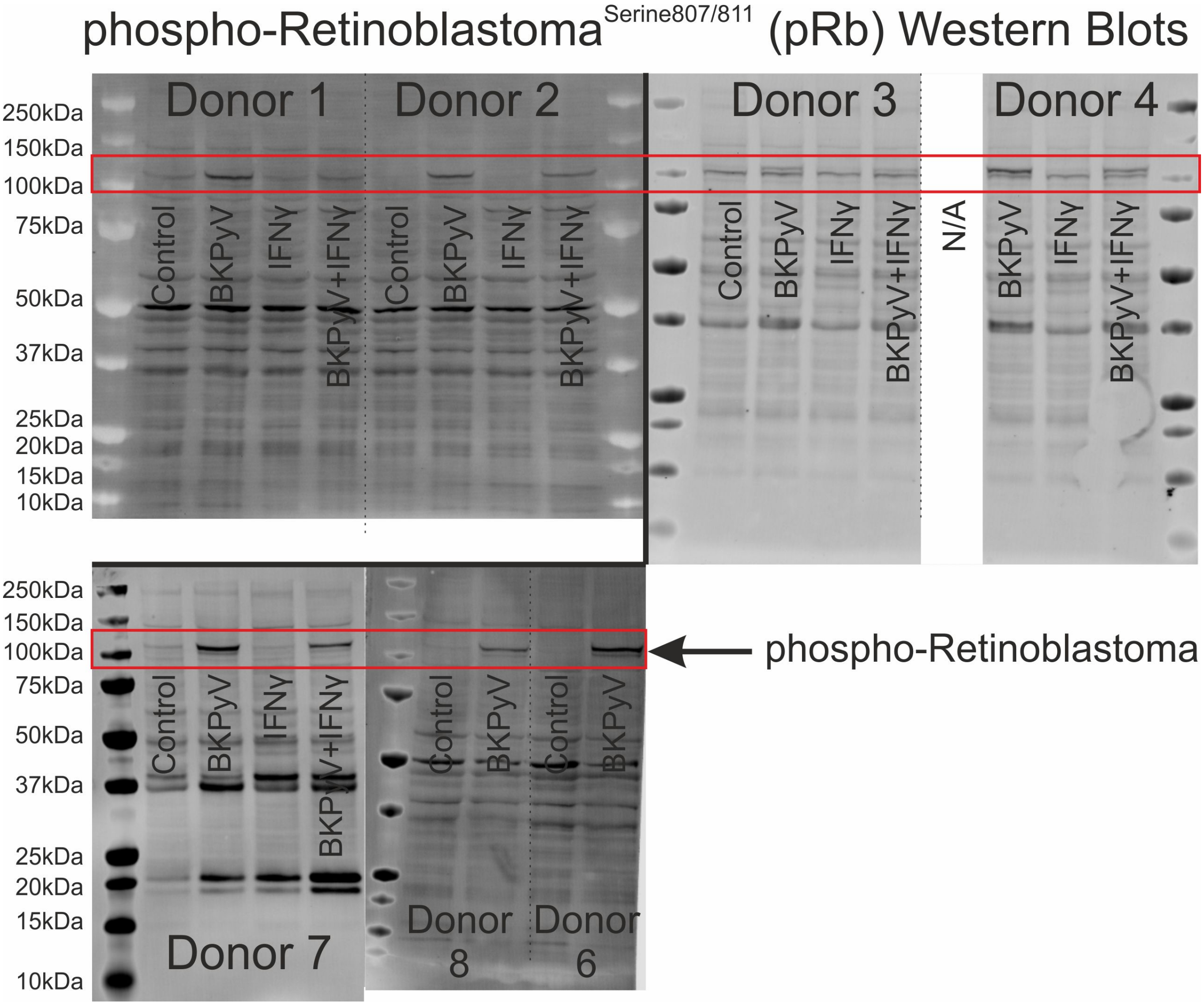


Extended Data Fig. 13 – Full anti-phosphorylated-Retinoblastoma (p-pRb) Western blots used for densitometry in Fig. 4. Predicted molecular weight for pRb is 106.2kDa (UniProtKB – P06400). This antibody recognises Retinoblastoma when phosphorylated at serine 807 or serine 811 and this is reflected in the presence of a doublet band. The lower band reflects single phosphorylation at serine 807/811 and the higher band represents phosphorylation at both sites. The control cells for Donor 4 were lost to an infection during culture and were therefore not available (N/A) for analysis.

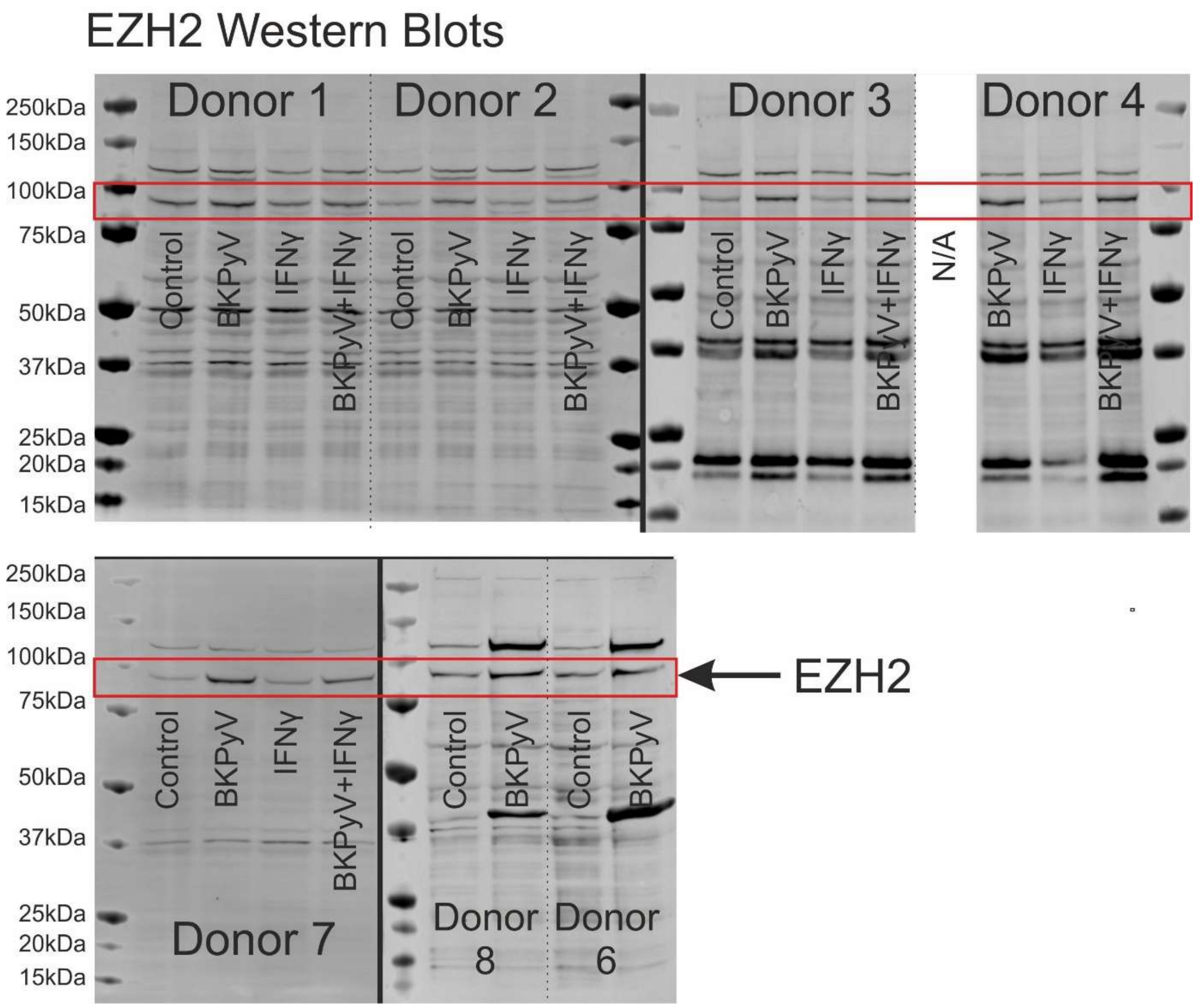


Extended Data Fig. 14 – Full EZH2 Western blots used for densitometry in Fig. 4. Predicted molecular weight for EZH2 is 85.4kDa (UniProtKB – Q15910). The control cells for Donor 4 were lost to an infection during culture and were therefore not available (N/A) for analysis.

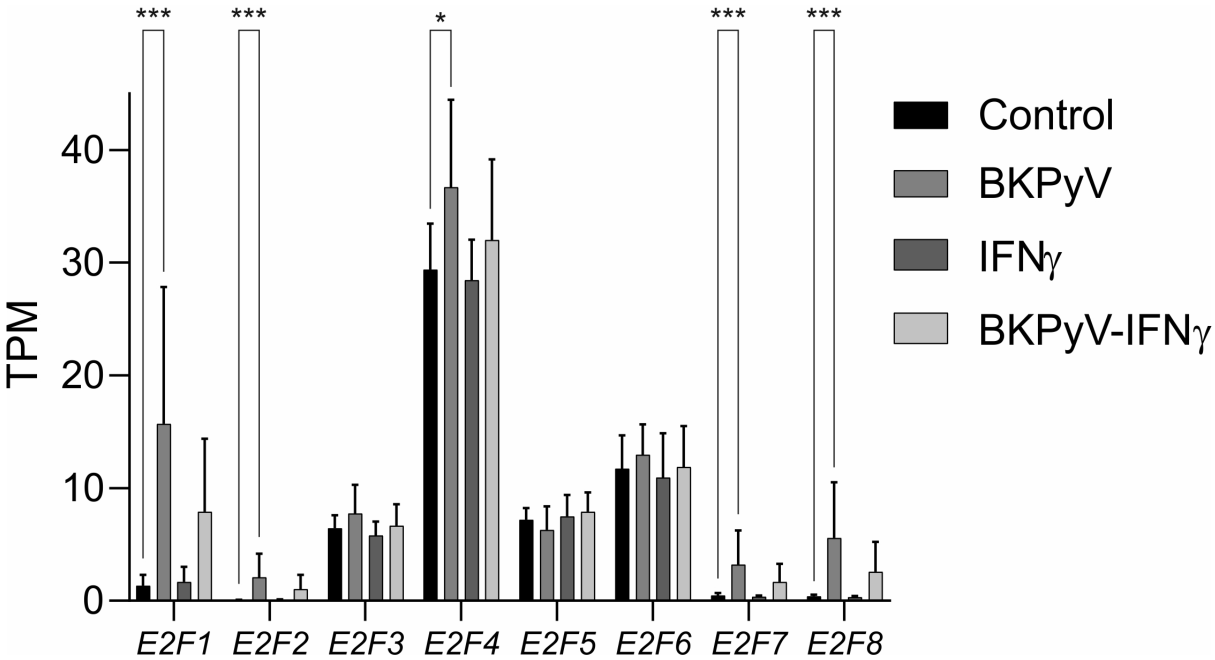


Extended Data Fig. 15 – mRNAseq data for the E2F family of transcription factors. The *E2F1* transcription showed the greatest fold change (mean $\log_2$ fold change (TPM+1) = 2.40; p<0.001). However, *E2F2* (mean $\log_2$ fold change (TPM+1) = 1.03; p<0.001), *E2F4* (mean $\log_2$ fold change (TPM+1) = 0.30; p=0.014), *E2F7* (mean $\log_2$ fold change (TPM+1) = 1.07; p<0.001) and *E2F8* (mean $\log_2$ fold change (TPM+1) = 1.64; p<0.001) were all induced significantly too.

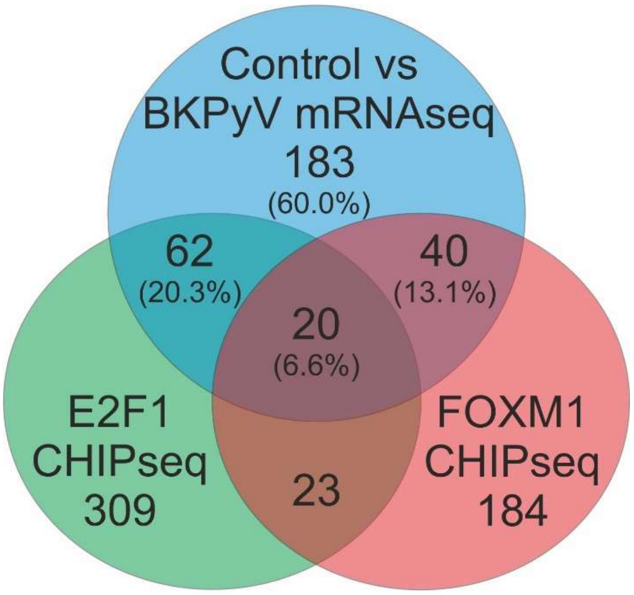


BKPyV - E2F1 CHIPseq Shared

| | |
|---|---|
| *ASF1B* | ***MCM2*** |
| *BLM* | *MCM3* |
| *BUB1B* | *MCM4* |
| *C1orf112* | *MCM6* |
| *C21orf58* | *MCM7* |
| *CDC25A* | *NCAPG* |
| *CDC25C* | *NCAPG2* |
| *CDC6* | *NDC80* |
| *CDC7* | *OIP5* |
| *CDCA3* | *PAQR4* |
| *CDCA5* | *PCNA* |
| *CENPK* | *PRR11* |
| *CENPM* | ***RAD51AP1*** |
| *CHAF1A* | *RAD54L* |
| *CKS1B* | *RECQL4* |
| *CLSPN* | *RFC4* |
| *DDX11* | *RRM1* |
| *DERL3* | *SKA1* |
| *DNAJC9* | *SLFN13* |
| *DONSON* | *SMC4* |
| *DSN1* | *TCF19* |
| *DTL* | *TIPIN* |
| *ESCO2* | *TUBA1B* |
| *EXO1* | *TYMS* |
| ***EZH2*** | *UBE2T* |
| *FAM111B* | *UHRF1* |
| *FANCI* | *WDR62* |
| *GTSE1* | *WDR76* |
| *HAUS8* | *XRCC2* |
| *MAD2L1* | *ZNF367* |
| *MASTL* | *ZWINT* |

BKPyV - FOXM1 CHIPseq Shared

| | |
|---|---|
| *ARHGAP11A* | *FBXO5* |
| *ASPM* | *GAS2L3* |
| *AURKA* | *HMMR* |
| *BRIP1* | *KIF18B* |
| *CCNA2* | *KIF20A* |
| *CCNB1* | *KIF20B* |
| *CCNB2* | *KIF23* |
| *CDC20* | ***MKI67*** |
| *CDC25B* | *NCAPH* |
| *CDCA2* | *NEK2* |
| *CDCA8* | *PHF19* |
| *CDKN3* | *PLK1* |
| *CENPF* | *PTTG1* |
| *CEP55* | *RACGAP1* |
| *CIT* | *SPC25* |
| *CKAP2L* | *TOP2A* |
| *DEPDC1* | *TPX2* |
| *DLGAP5* | *TROAP* |
| *ESPL1* | *TTK* |
| *FAM83D* | *UBE2C* |

Shared by all

| | |
|---|---|
| *ATAD2* | *KIFC1* |
| *AURKB* | *LMNB1* |
| *CDK1* | *MXD3* |
| *CENPA* | *NUF2* |
| *DEPDC1B* | *NUSAP1* |
| *HJURP* | *PRC1* |
| *HMGB2* | *RRM2* |
| *KIF11* | *SPAG5* |
| *KIF18A* | *TMPO* |
| *KIF2C* | *UBE2S* |

Extended Data Fig. 16 – Comparison of genes significantly induced by >2-fold following BKPyV infection with published CHIPseq peaks for E2F1 in MM1.S cells [2] and FOXM1 in U20S cells [3] reveals significant overlap. E2F1 and FOXM1 CHIPseq peaks were found near 33.6% and 29.5% of BKPyV-induced genes, respectively. The significance of the overlap was tested by calculating their exact hypergeometric probability. For FOXM1: representation factor = 40.3 and $p < 2.70e-78$. For E2F1: representation factor = 35.6 and $p < 9.88e-103$.

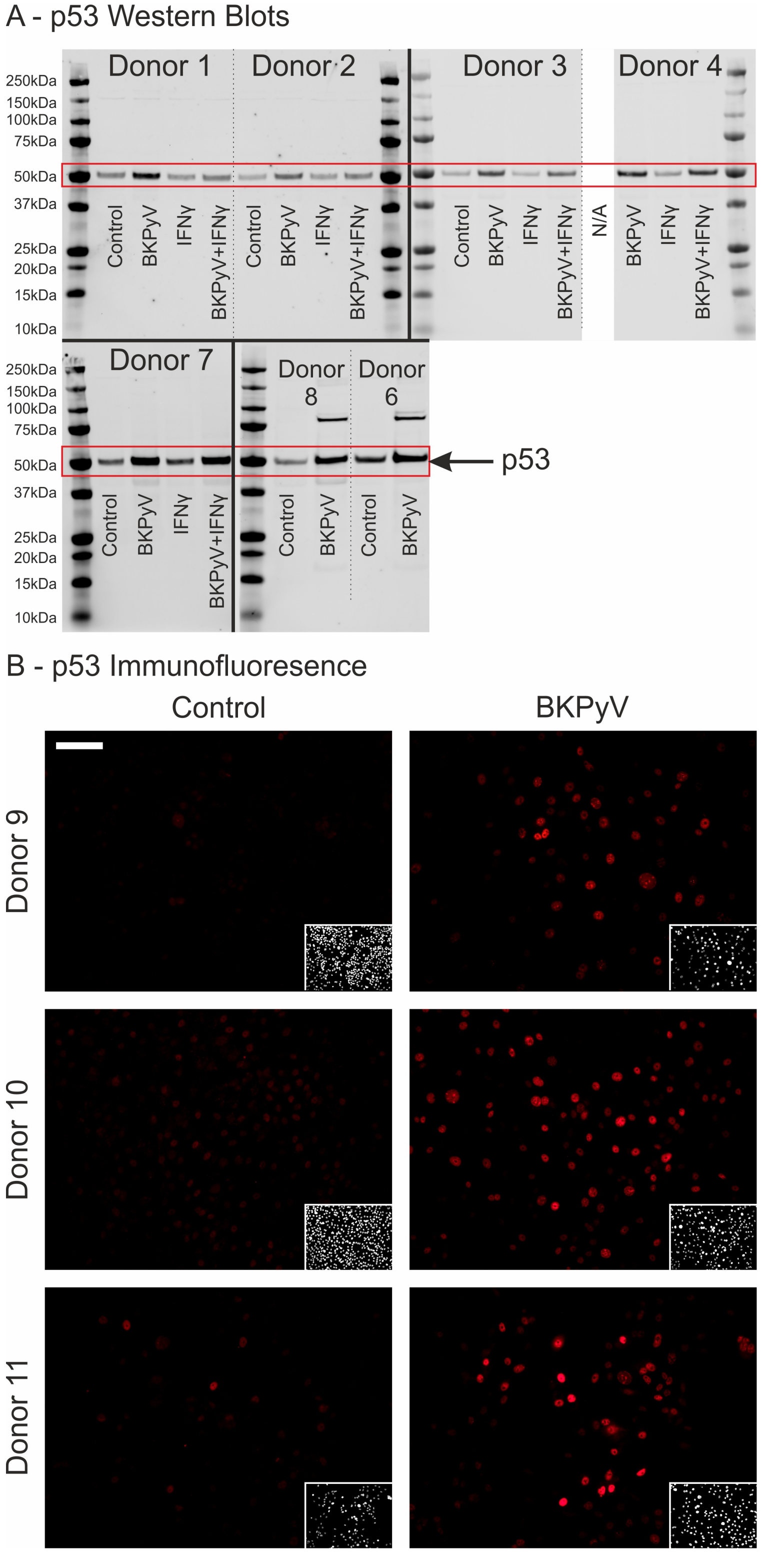


Extended Data Fig. 17 – (A) Full p53 Western blots used for densitometry in Fig. 5. Predicted molecular weight for p53 is 53kDa (UniProtKB – P04637). The higher bands on the blots for Donor 8 and 6 reflect a previous probing of the membrane for LT-Ag (see Extended Data Fig. 3). The control cells for Donor 4 were lost to an infection during culture and were therefore not available (N/A) for analysis.

(B) Indirect immunofluorescence for p53 found stabilised p53 protein was localised to the nuclei of BKPyV infected urothelial cells (n=3 independent donors). Scale bar in Donor 9 control main panel indicates 100 μm.

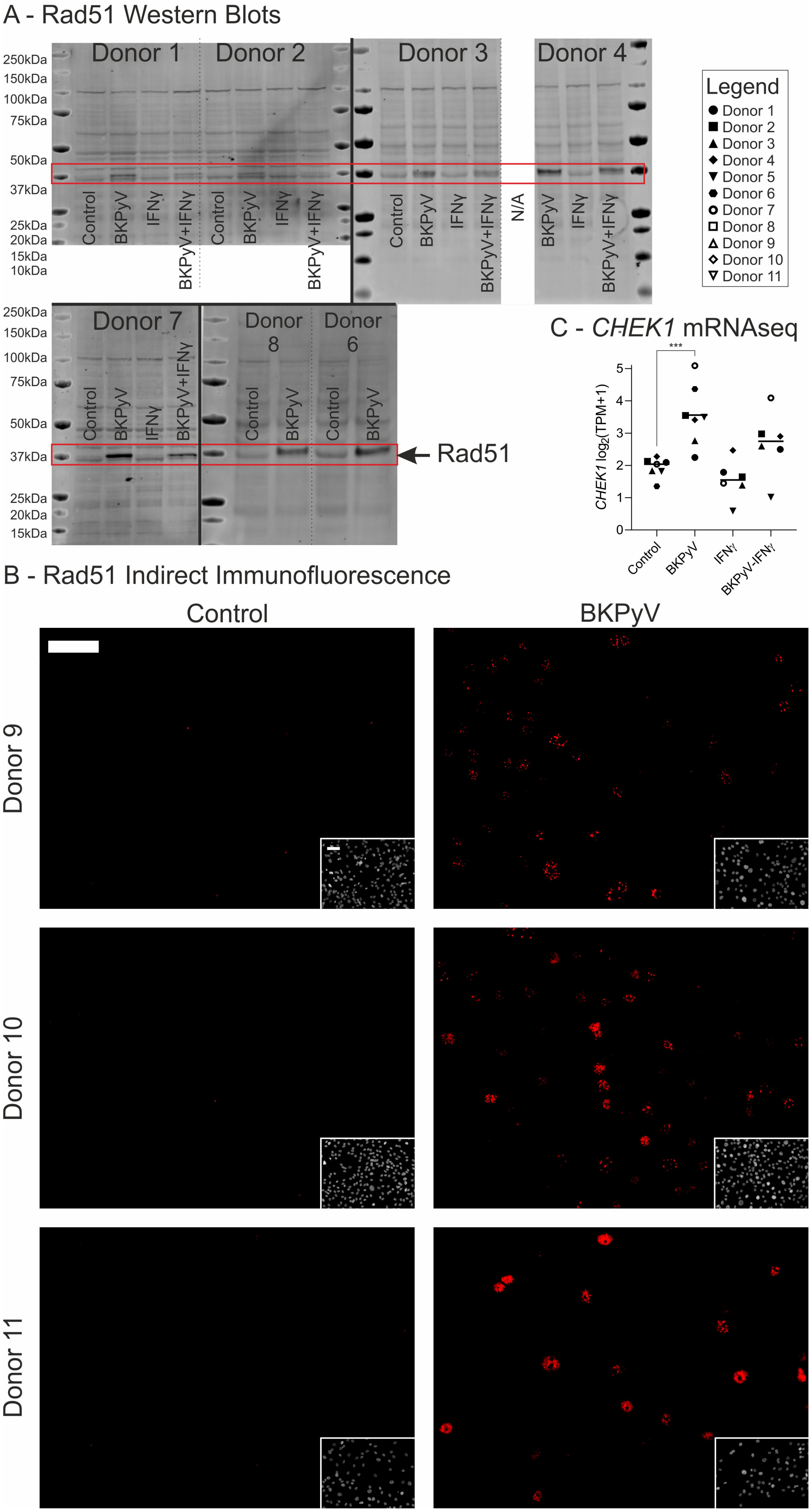
A - Rad51 Western Blots
Donor 1
Donor 2
Donor 3
Donor 4
250kDa
150kDa
100kDa
75kDa
50kDa
37kDa
25kDa
20kDa
15kDa
10kDa
Control
BKPyV
IFNγ
BKPyV+IFNγ
N/A
Legend
Donor 1
Donor 2
Donor 3
Donor 4
Donor 5
Donor 6
Donor 7
Donor 8
Donor 9
Donor 10
Donor 11
Donor 7
Donor 8
Donor 6
Rad51
C - CHEK1 mRNAseq
***
CHEK1 log2(TPM+1)
Control
BKPyV
IFNγ
BKPyV-IFNγ
B - Rad51 Indirect Immunofluorescence
Control
BKPyV
Donor 9
Donor 10
Donor 11

148

Extended Data Fig. 18 – (A) Full RAD51 Western blots used for densitometry in Fig. 5. Predicted molecular weight for RAD51 is 37.0kDa (UniProtKB – Q06609). The slight increase in molecular weight observed from control to BKPyV-infected lysates likely reflects phosphorylation of the Rad51 protein, which has previously been associated with activity. The control cells for Donor 4 were lost to an infection during culture and were therefore not available (N/A) for analysis. (B) Indirect Immunofluoresence for Rad51 showed speckles within the nuclei of BKPyV infected urothelial cells (n=3 independent donors). Scale bars in the top left corner of donor 9 Control main image and inset Hoescht 33258 stain both denote 100μm. (C) Chk1 kinase performs activating phosphorylation of Rad51. Expression of the *CHEK1* gene was significantly ($p$=0.0001) induced by BKPyV infection.

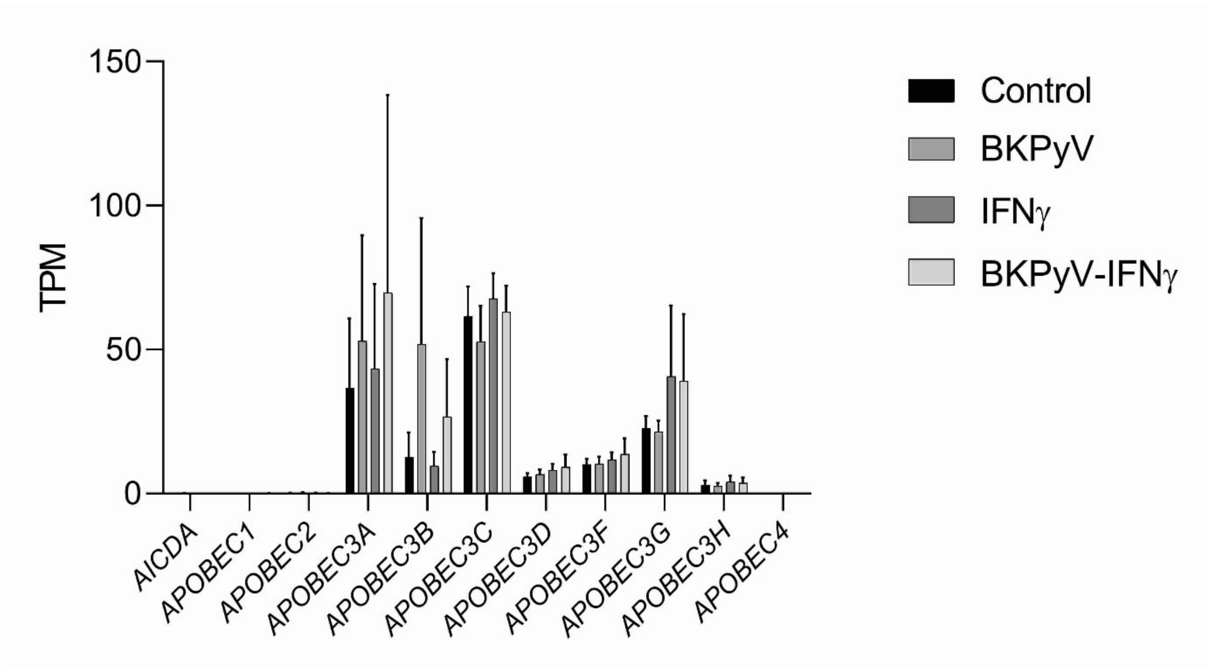


Extended Data Fig. 19 – Analysis of cytosine deaminase gene transcription by cultures of differentiated normal human urothelial (NHU) cells (cell lines developed from n=7 independent donors, positive error bars denote standard deviation).

A

| | *st-Ag* | *LT-Ag* | *Agnoprotein* | *VP1* | *VP2* |
|---|---|---|---|---|---|
| *APOBEC3A* | 0.32 ($p$=0.283) | 0.39 ($p$=0.185) | 0.41 ($p$=0.157) | 0.11 ($p$=0.729) | 0.28 ($p$=0.354) |
| *APOBEC3B* | 0.97 ($p$=1.26x10$^{-8}$) | 0.98 ($p$= 1.41x10$^{-9}$) | 0.95 ($p$=1.55x10$^{-7}$) | 0.81 ($p$=4.51x10$^{-4}$) | 0.95 ($p$=1.93x10$^{-7}$) |

B

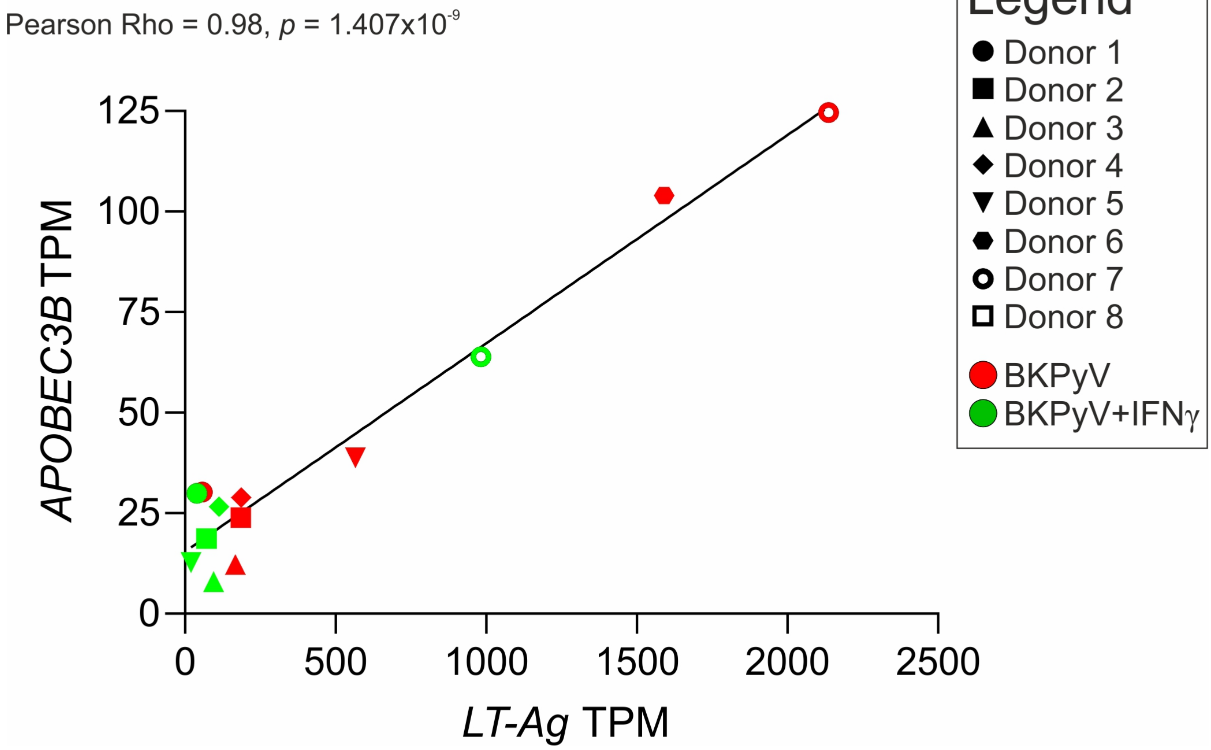


Extended Data Fig. 20 – (A) Table of Pearson correlation Rho values (and significance *p* values) for comparison of *APOBEC3A* and *APOBEC3B* TPMs with viral transcript relative TPMs. (B) Linear regression analysis of the relationship between BKPyV *LT-Ag* TPM and *APOBEC3B* TPM.

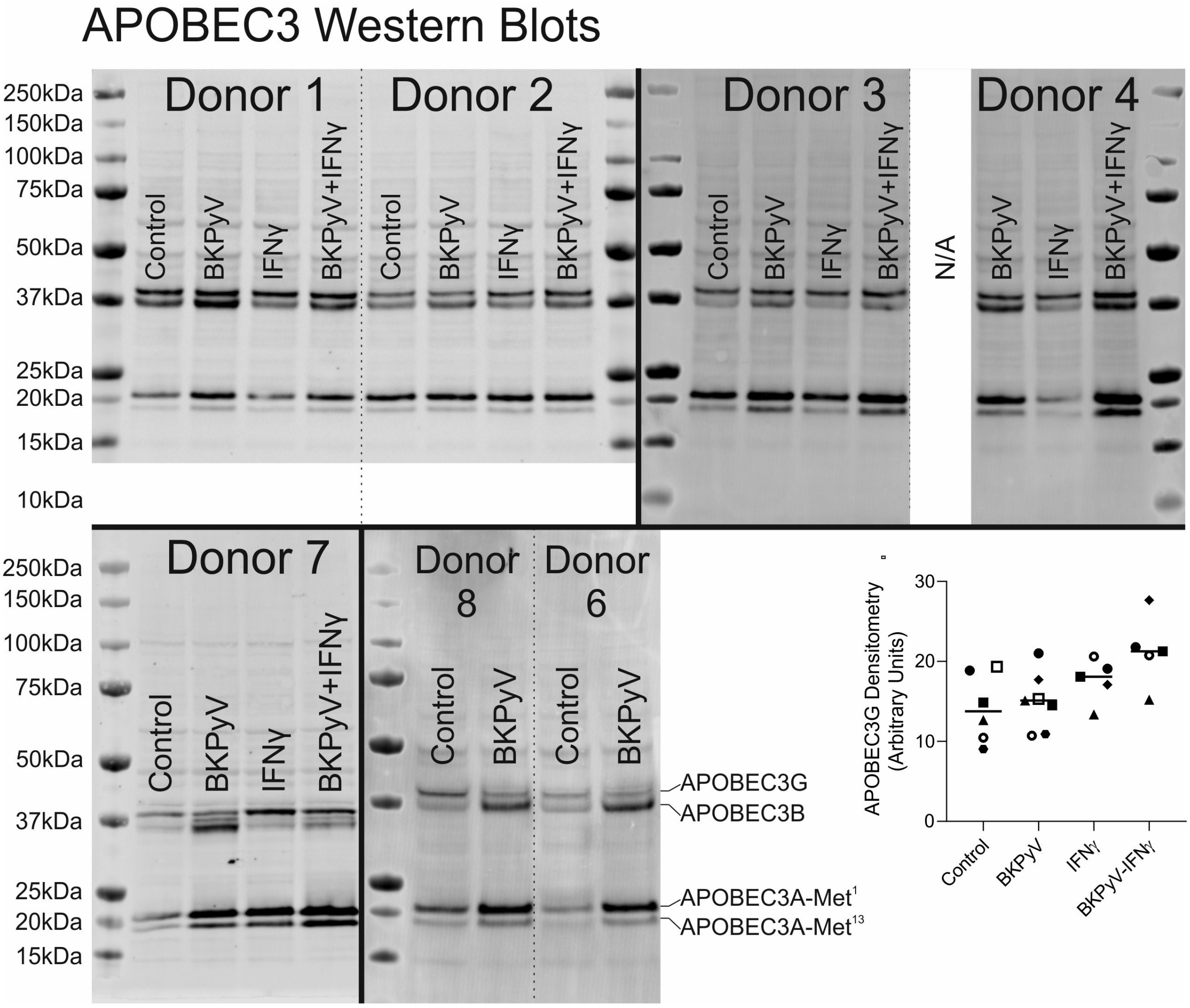


Extended Data Fig. 21 – Full anti-APOBEC3 Western blots used for densitometry in Fig. 6. The anti-APOBEC3A/B/G rabbit monoclonal (clone 5210-87-13) antibody was developed in the laboratory of Reuben Harris [4]. A detailed characterisation demonstrating the reactivity of this antibody against all APOBEC3 isoforms was recently published showing it detects APOBEC3A, APOBEC3B and APOBEC3G which are readily distinguished by molecular weight [15]. The predicted molecular weight for APOBEC3A (UniProtKB – P31941) is 23kDa ($Met^{1}$) and a smaller 21.8kDa enzymatically-active variant is generated by internal translation initiation at a methionine at position 13 ($Met^{13}$) [5]. The predicted molecular weight for APOBEC3B is 45.9kDa (UniProtKB - Q9UH17). The predicted molecular weight for APOBEC3G is 46.4kDa (UniProtKB - Q9HC16). However, the running of APOBEC3B and APOBEC3G around the 37kDa marker in Western blots is consistent with previous reports [6]. The control cells for Donor 4 were lost to an infection during culture and were therefore not available (N/A) for analysis.

Densitometry analysis for the APOBEC3G band is also included here but showed no significant changes in spite of apparent IFNγ-mediated induction at the transcript level (Extended Data Fig. 19).

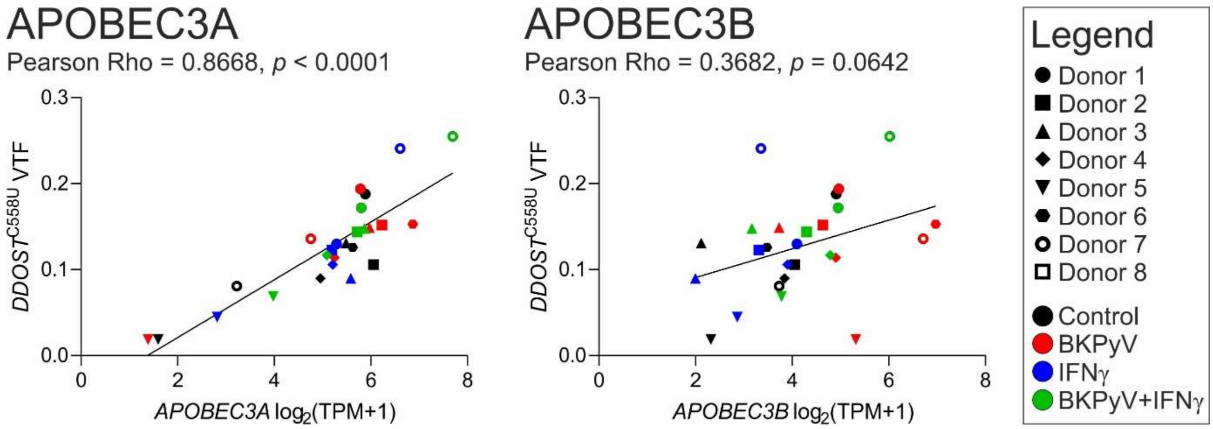


Extended Data Fig. 22 – Correlation analysis of the relationship between APOBEC3 enzyme transcript expression and DDOST C558U variant transcript abundance.

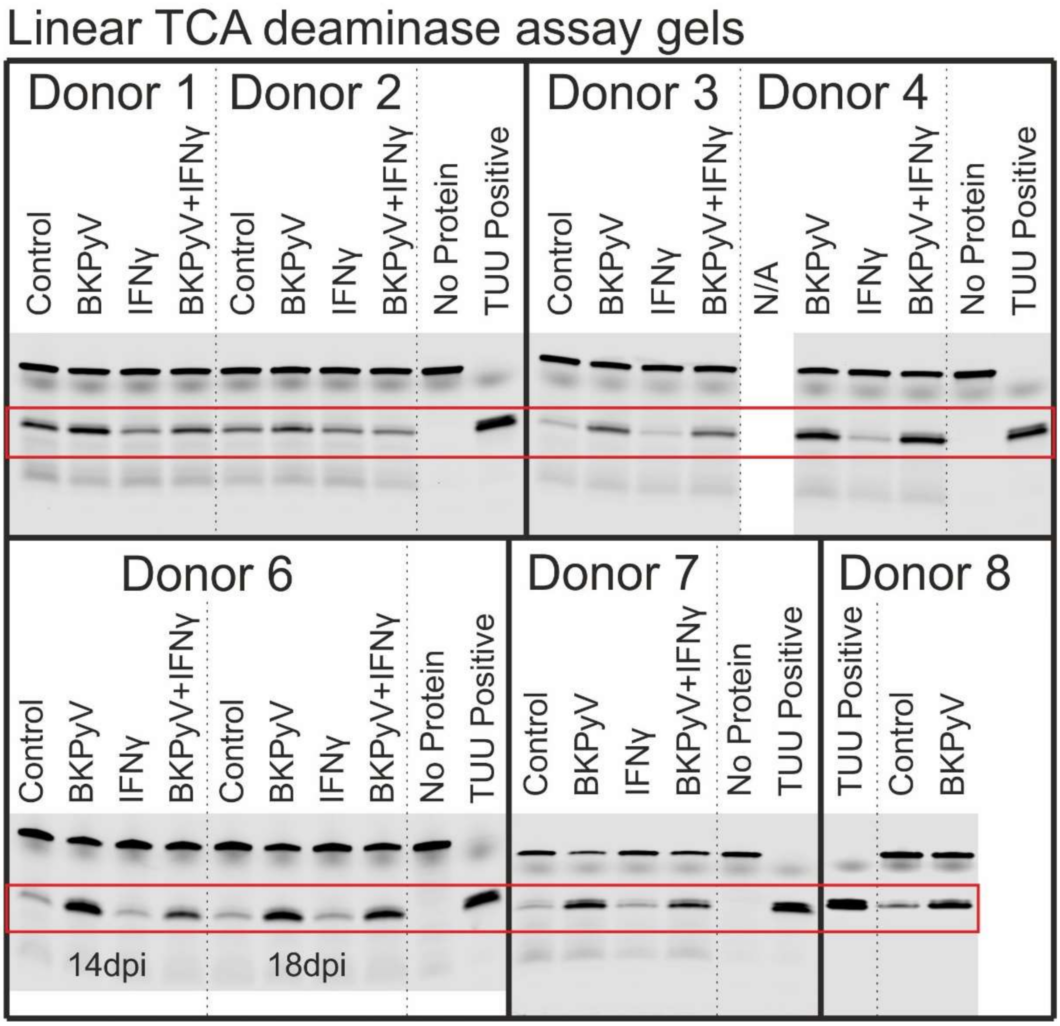


Extended Data Fig. 23 – Deaminase assays performed with a linear RTCA probe preferential for APOBEC3B. The red box shows the band for the cleaved probe which was analysed by densitometry for panel Fig. 6F. The Donor 6 18dpi assays were not included in the densitometry shown in Fig. 6.

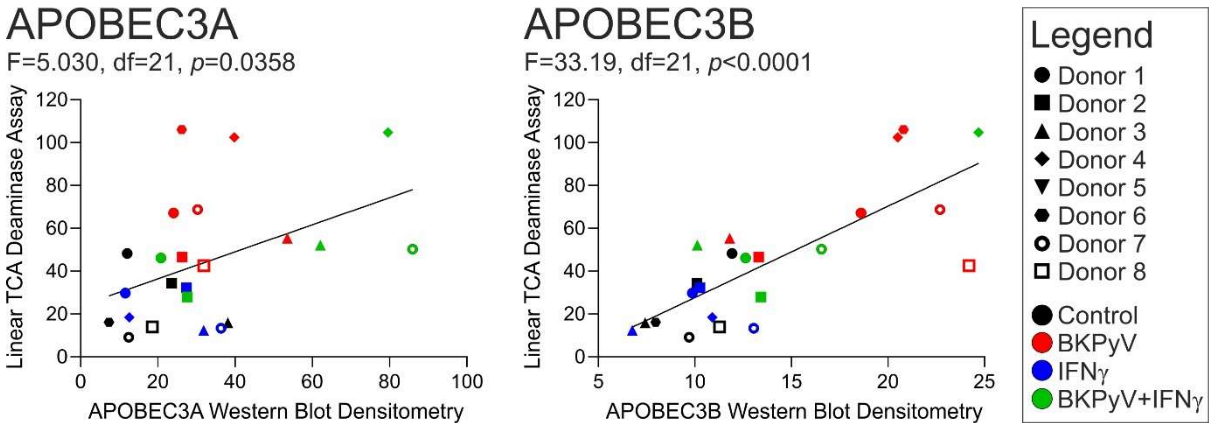


Extended Data Fig. 24 – Linear regression analysis of the relationship between linear TCA deaminase assays and APOBEC3A and APOBEC3B protein abundance (assessed by Western blot in Fig. 6).

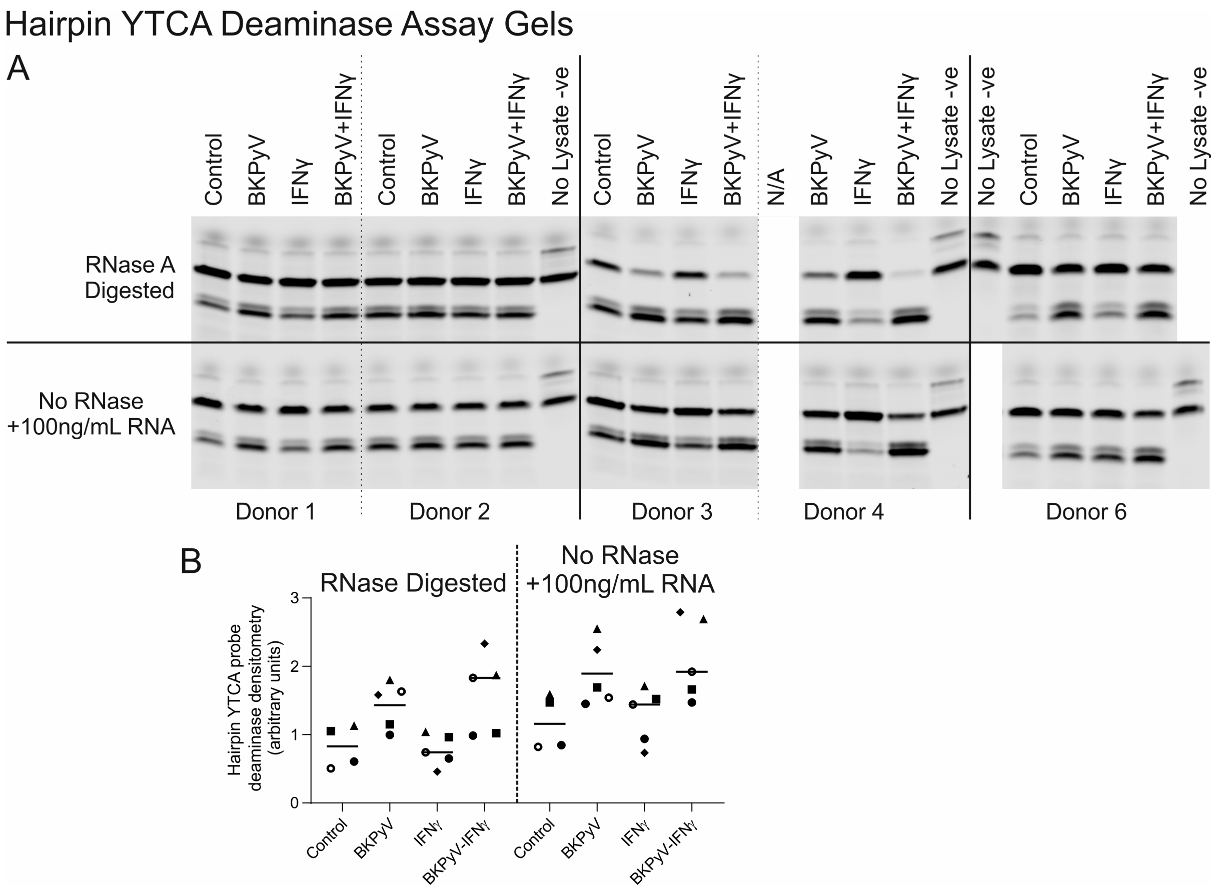


Extended Data Fig. 25 – (A) Full gel scans for deaminase assays performed with a hairpin YTCA probe previously reported to be preferential for APOBEC3A in the presence of exogenous RNA [7]. The presence of RNA was previously shown to inhibit APOBEC3B giving this assay APOBEC3A specificity [7]. (B) Densitometry for the deaminase assays performed with a hairpin YTCA probe, found the addition of exogenous RNA had no effect on deaminase activity in BKPyV-infected urothelial cell lysates.

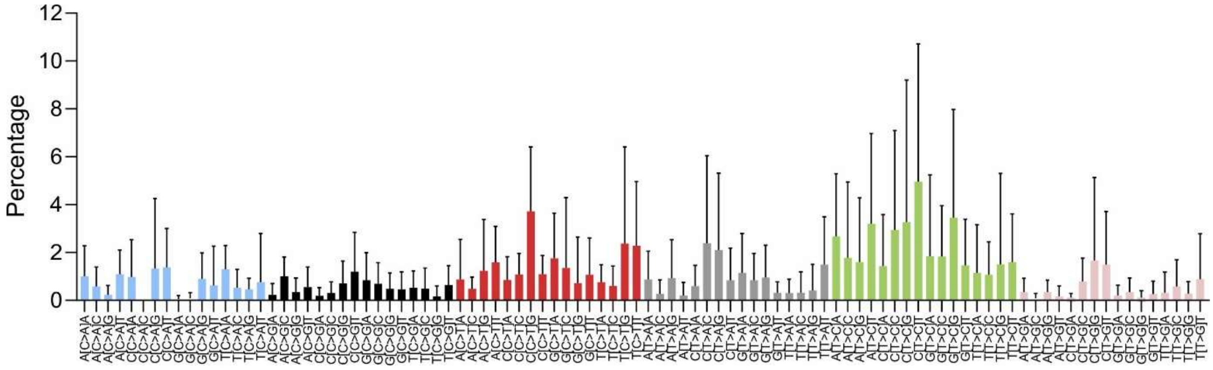


Extended Data Fig. 26 – Single base substitution mRNAseq mutational signature induced by BKPyV-infection.

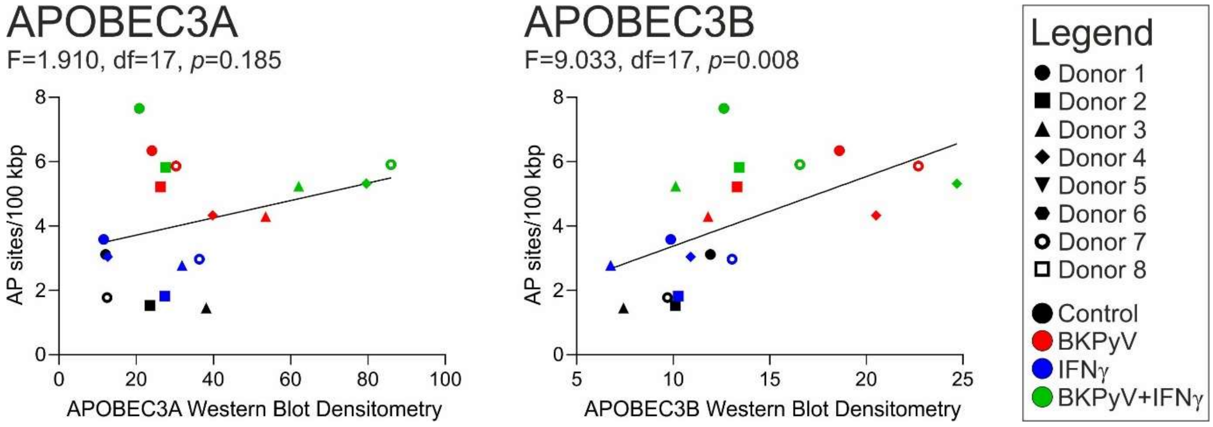


Extended Data Fig. 27 – Linear regression analysis of the relationship between the formation of apurinic/apyrimidinic (AP) sites in the host urothelial genome and APOBEC3A and APOBEC3B protein abundance (assessed by Western blot in Fig. 6).

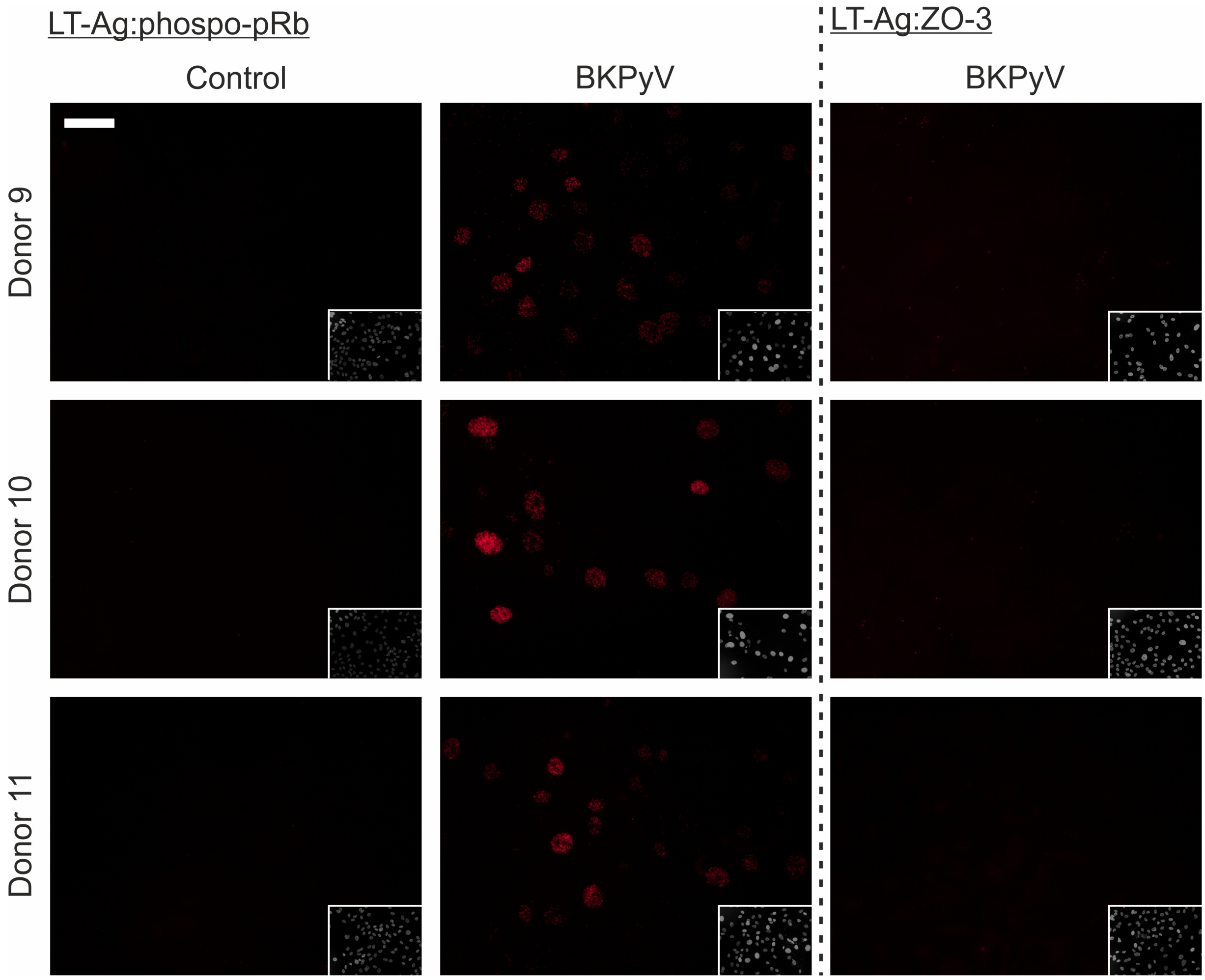


Extended Data Fig. 28 – Proximity ligation controls.

Direct binding of Large T antigen (LT-Ag) and Retinoblastoma protein (pRb) has previously been reported[8]. The LT-Ag:pRb interaction was therefore included as a positive control for the assay. LT-Ag interactions were always negative in uninfected controls.

Zonula Occludins 3 is a tight junction protein that localises to the plasma membrane and was therefore unlikely to interact with the predominantly nuclear LT-Ag. LT-Ag:ZO-3 was included as a negative control antibody pair.

Images of DAPI stained nuclei are inset. White scale bar in the main panel of Donor 9 Control indicates 50 μm.

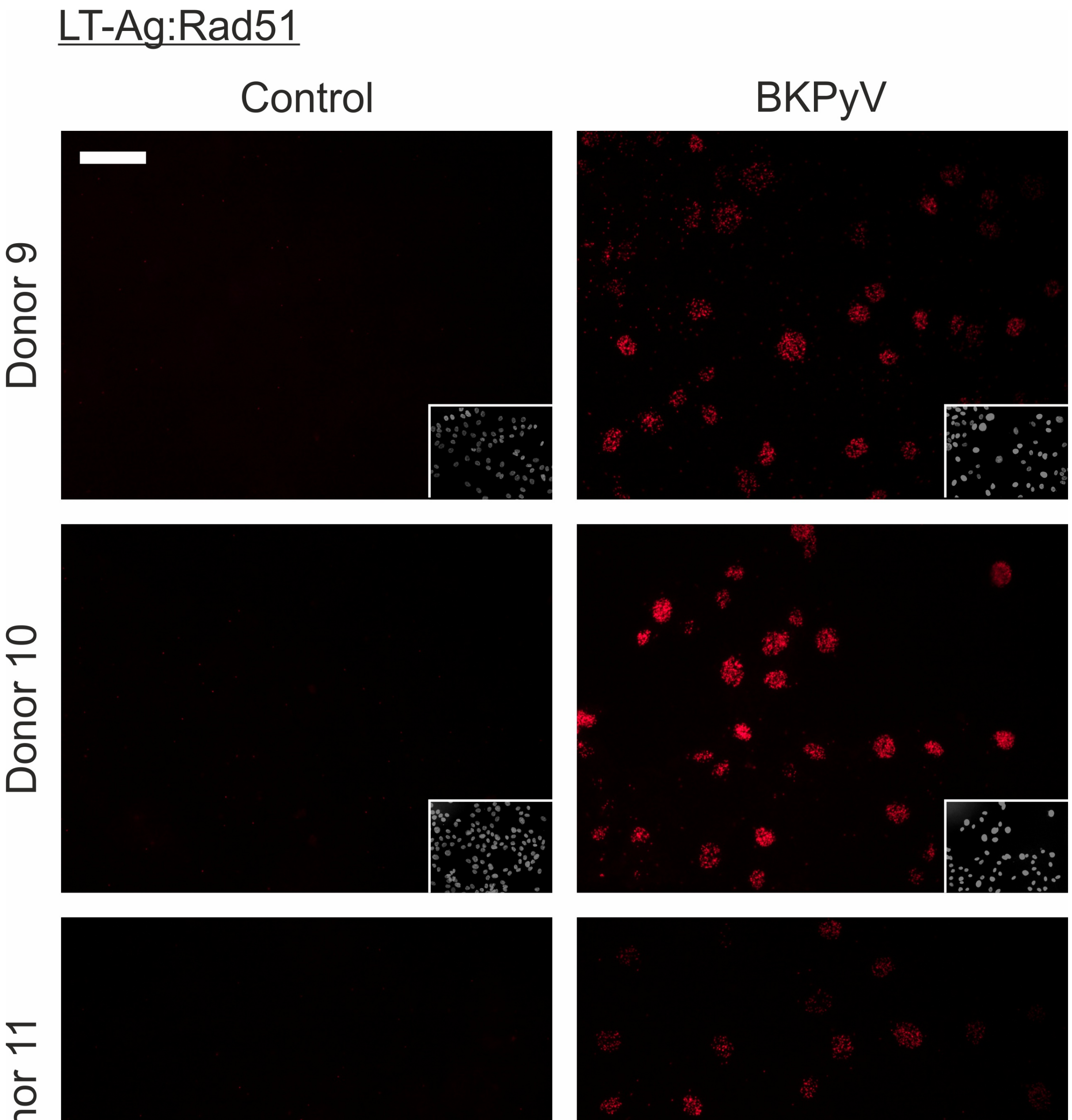


Extended Data Fig. 29 – Proximity ligation of Large T antigen (LT-Ag) and Rad51 was negative in non-infected control normal human urothelial cells but showed positive nuclear speckles in BKPyV-infected cells. Images of DAPI stained nuclei are inset. White scale bar in the main panel of Donor 9 Control indicates 50 μm.

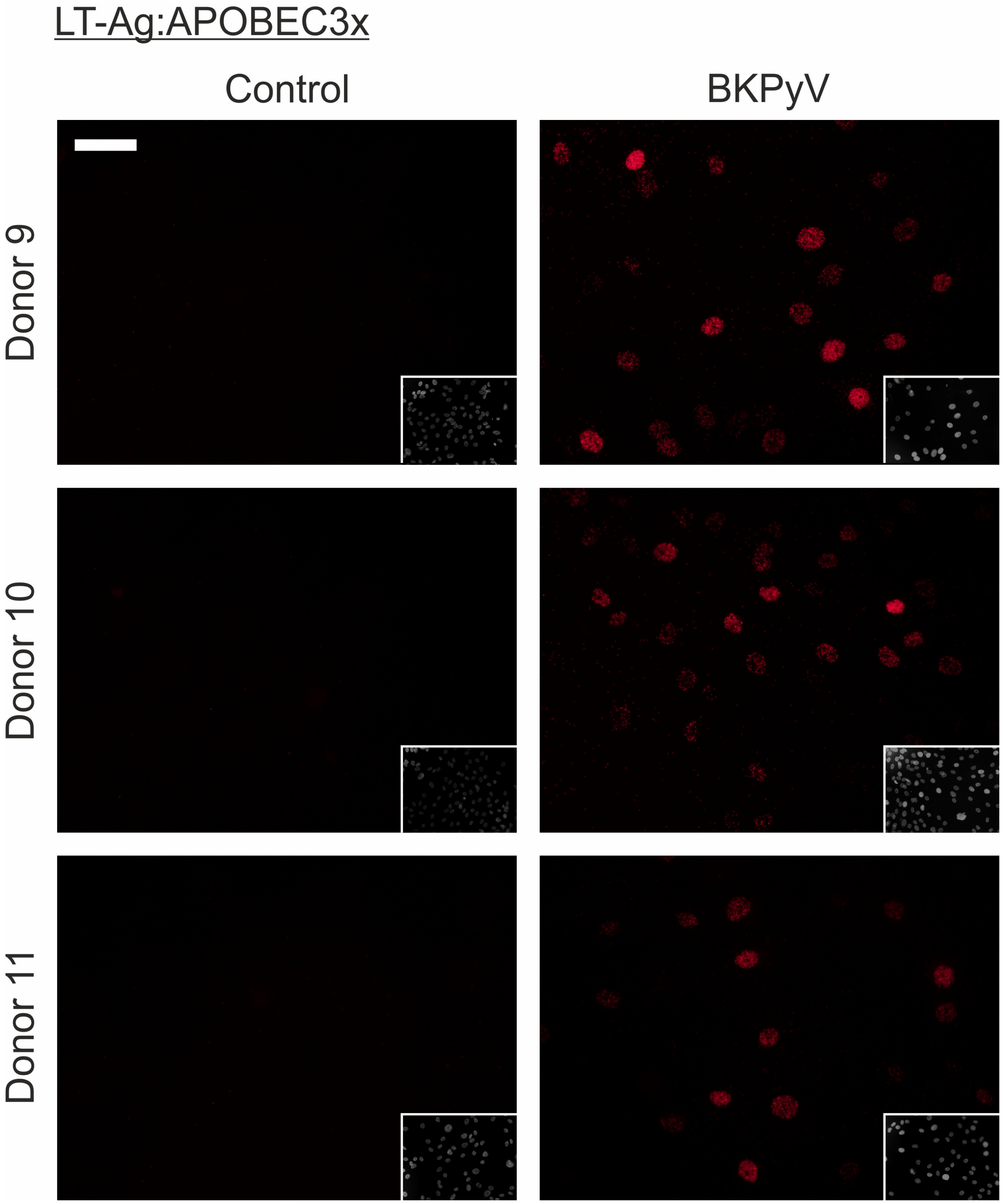


Extended Data Fig. 30 – Proximity ligation of Large T antigen (LT-Ag) and the APOBEC3A/B/G antibody[4] was negative in non-infected control normal human urothelial cells but showed positive nuclear speckles in BKPyV-infected cells. Images of DAPI stained nuclei are inset. White scale bar in the main panel of Donor 9 Control indicates 50 μm.

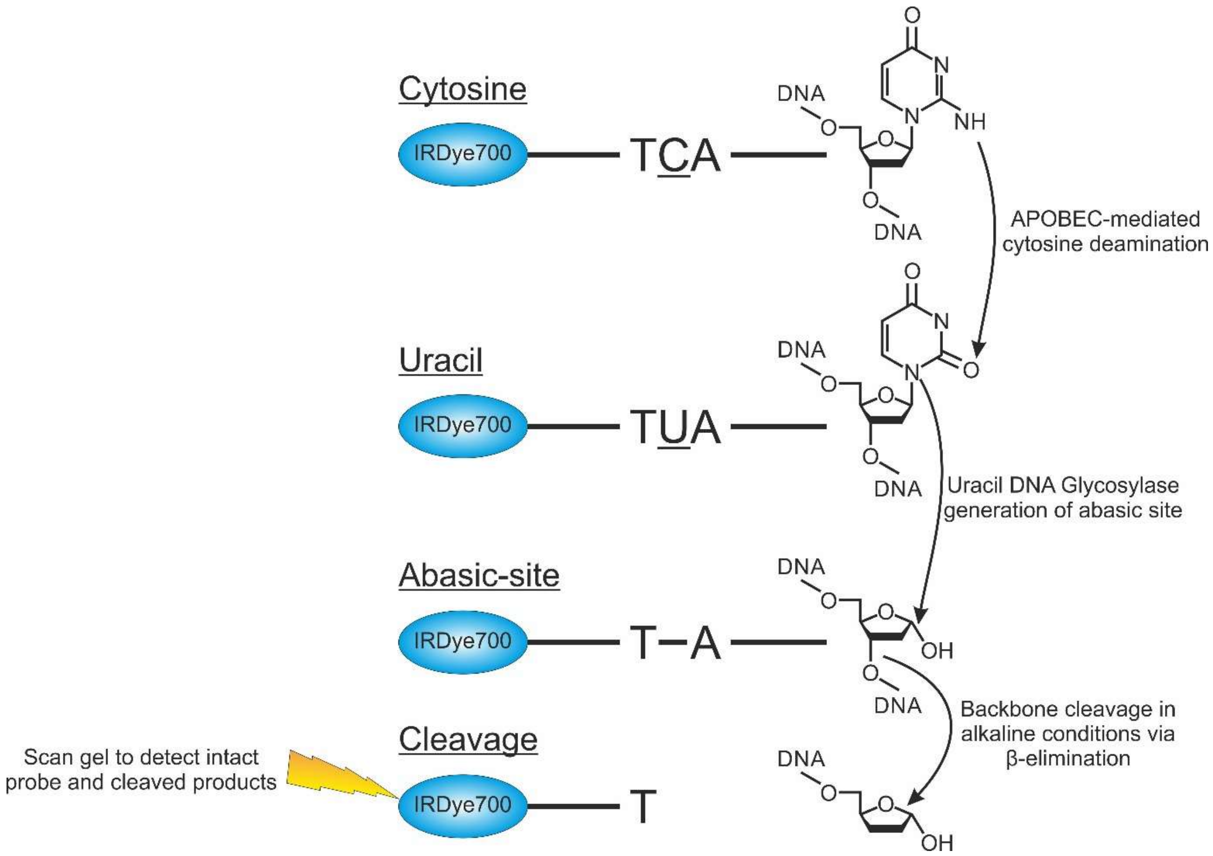


Extended Data Fig. 31 – Schematic describing the deaminase assay method principle. Fluorescently labelled ssDNA probes containing either a RTCA or YTCA motif were exposed to the APOBEC enzymes present in NHU cell lysates. APOBECs deaminate the cytosine bases in a TCA context to uracil. Exposure to uracil DNA glycosylase (UDG) cleaves off the uracil leaving an abasic site. The reaction buffer is then turned alkaline by the addition of NaOH which leads to cleavage of the DNA backbone by β–elimination. The intact probe and the cleaved DNA attached to the probe can then be run on a gel and detected.
